# Mechanics regulate human embryonic stem cell self-organization to specify mesoderm

**DOI:** 10.1101/2020.02.10.943076

**Authors:** Jonathon M. Muncie, Nadia M.E. Ayad, Johnathon N. Lakins, Valerie M. Weaver

## Abstract

Embryogenesis is directed by morphogens that induce differentiation within a defined tissue geometry. Tissue organization is mediated by cell-cell and cell-extracellular matrix (ECM) adhesions and is modulated by cell tension and tissue-level force. Whether cell tension regulates development by directly influencing morphogen signaling remains unclear. Human embryonic stem cells (hESCs) exhibit an intrinsic capacity for self-organization that motivates their use as a tractable model of early human embryogenesis. We engineered patterned substrates that enhance cell-cell interactions to direct the self-organization of cultured hESCs into “gastrulation-like” nodes. Tissue geometries that generate local nodes of high cell-cell tension and induce these self-organized tissue nodes drive BMP4-dependent gastrulation by enhancing phosphorylation and nuclear translocation of β-catenin to promote Wnt signaling and mesoderm specification. The findings underscore the interplay between tissue organization, cell tension, and morphogen-dependent differentiation, and demonstrate that cell- and tissue-level forces directly regulate cell fate specification in early human development.

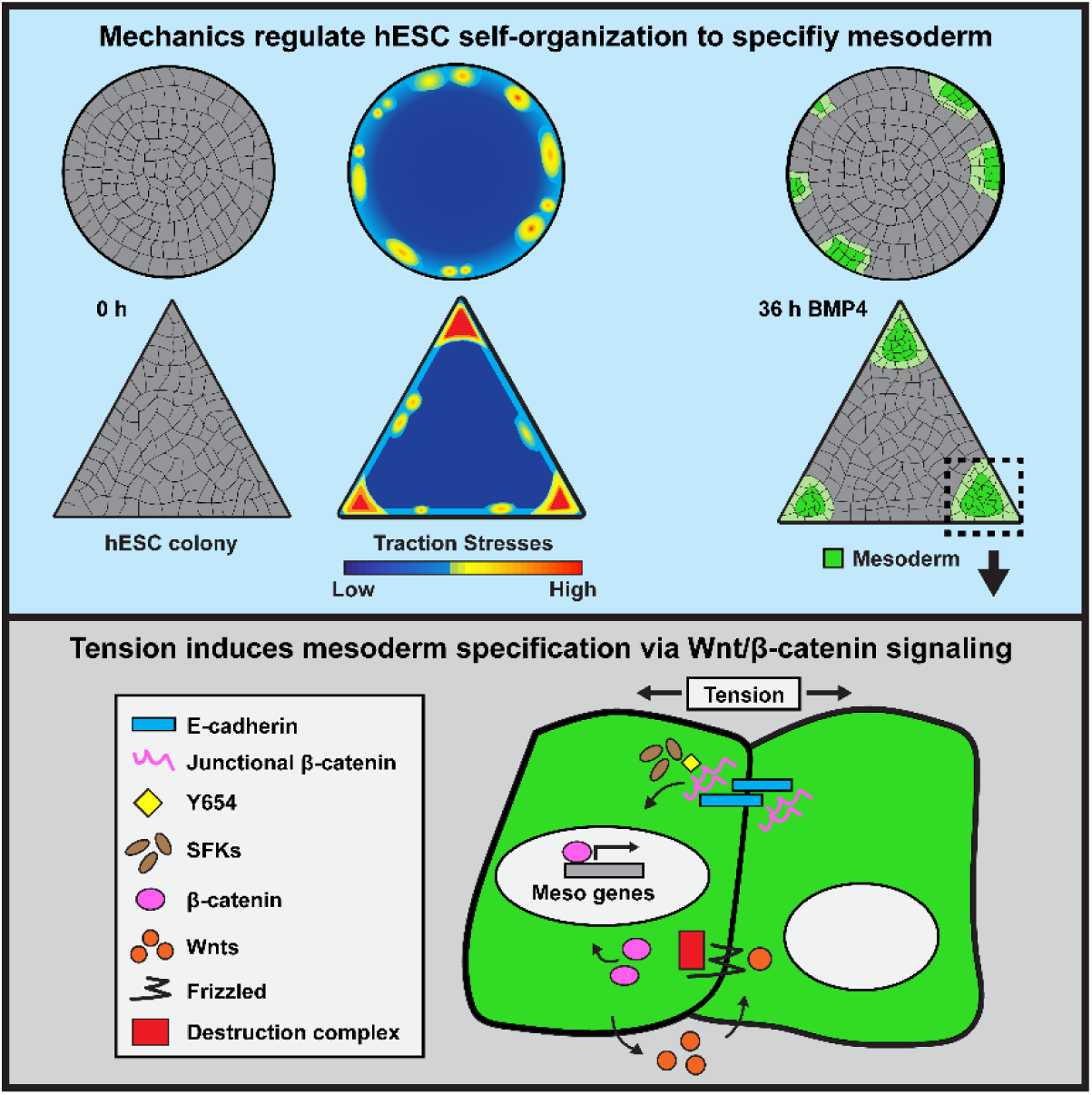
Graphical Abstract.

**Highlights:**

- Substrates that enhance cell-cell adhesion promote hESC self-organization
- Tissue nodes exhibiting high tension are predisposed to gastrulation induction
- Colony geometry dictates the localization of tension nodes to specify mesoderm
- Tension activates β-catenin and stimulates Wnt signaling to induce mesoderm

**In Brief:** Engineered substrates that promote cell-cell adhesion and reconstitute epiblast tissue organization facilitate “gastrulation-like” morphogenesis in cultured hESCs. Tissue geometries that foster localized regions of high cell-cell tension potentiate BMP4-dependent mesoderm specification by enhancing phosphorylation and nuclear translocation of β-catenin to promote Wnt signaling.

## Introduction

Early embryogenesis is highly complex and involves the spatiotemporal regulation of cell differentiation that is critical for driving tissue development and guiding morphogenesis. Gastrulation in particular, requires precise coordination of cell fate specification, wherein the cells of the epiblast simultaneously segregate and differentiate into the three primary germ layers: ectoderm, mesoderm, and endoderm. Gastrulation is stimulated by morphogens and growth factors, modulated by their respective inhibitors, and directed by transcriptional programs that are influenced by epigenetics (Marlow, 2019; Molè et al., 2020; Raffaelli and Stern, 2020; Stathopoulos and Newcomb, 2020). The morphogenesis that occurs during gastrulation relies on cell tension and tissue-level force to facilitate the physical segregation of epiblast cells into the primary germ layers (Hamada, 2015; Ko and Martin, 2020; Paré and Zallen, 2020; Voiculescu et al., 2014; Williams and Solnica-Krezel, 2017). As embryogenesis progresses, tissue-level forces sculpt the embryo by driving neural tube closure, mediating somitogenesis, and guiding heart tube looping (McMillen and Holley, 2015; Ramasubramanian et al., 2013; Taber, 2014; Turlier and Maître, 2015; Vijayraghavan and Davidson, 2017). Thus, cell- and tissue-level force plays an intimate role in directing the structure-function of the emerging organism. Importantly, in the adult organism, cytoskeletal tension also regulates receptor activity and signaling, and modulates gene transcription to direct cell growth, survival, migration, and differentiation (Eliazer et al., 2019; McBeath et al., 2004; Paszek et al., 2005; Rys et al., 2015; Wang et al., 2012). Whether cytoskeletal tension could similarly influence developmental processes, such as gastrulation, by modifying cell signaling, transcription, and tissue-specific differentiation, and how, remains unclear.

Model organisms are powerful systems with which to study early development, including the impact of tissue-level force and cellular tension (Hiramatsu et al., 2013; Zhang et al., 2019). Nevertheless, direct quantitative measurements of cell tension and long-term high-resolution imaging are difficult to perform on intact embryos, presenting a major challenge for understanding how cell- and tissue-level forces regulate development. Remarkably, hESCs will self-organize to recapitulate patterns of the primary germ layers, even in the absence of extraembryonic tissue (Shao et al., 2017; Simunovic et al., 2019; Warmflash et al., 2014; Zheng et al., 2019). Prior studies using hESCs underscored the importance of tissue organization in early human tissue development, and implicated morphogen gradients and receptor accessibility as two key mechanisms whereby tissue geometry specifies primary germ layer differentiation (Blin et al., 2018; Chhabra et al., 2019; Etoc et al., 2016; Manfrin et al., 2019; Martyn et al., 2019; Smith et al., 2018; Tewary et al., 2019; Warmflash et al., 2014). Tissue-level force also increases in multicellular tissues, and cell-cell tension, in particular, is spatially enhanced by specific tissue geometries (Gomez et al., 2010; Kilian et al., 2010; Lee et al., 2016; Nelson et al., 2005). This raises the intriguing possibility that tissue geometry modulates local cytoskeletal tension to spatially direct hESC self-organization and tissue-specific differentiation.

Here we explored the role of cell-adhesion-mediated cytoskeletal tension in the generation of spatially-restricted, “gastrulation-like” nodes in hESCs, and monitored how this tension impacted mesoderm specification. We used hESCs cultured on biochemically-defined and geometrically-patterned ECMs of tuned elasticity to recapitulate the biophysical properties and promote the spatially-directed “gastrulation-like” tissue structure of the early embryo. Using this system, we implicated cytoskeletal tension as a key regulator of tissue self-organization and determined that cell-cell tension promotes mesoderm specification by modulating the activity of a key transcription factor that regulates Wnt signaling. The findings reveal, for the first time, a direct relationship between cell-cell tension and morphogen-dependent development, and emphasize the versatility of hESCs as a tractable model to study early human development.

## Results

### Compliant substrates promote hESC self-organization into “gastrulation-like” nodes

To study the interplay between tissue structure and early development, we designed reproducible culture strategies that foster hESC self-organization and consistently permit gastrulation induction following BMP4 stimulation. hESCs plated as confined circular colonies on ECM-patterned rigid glass substrates (Elastic modulus, E = 10^9^ to 10^10^ Pa) exhibit a radial pattern of primary germ layer differentiation following BMP4 stimulation (Warmflash et al., 2014). This has been attributed to restriction of BMP4/SMAD signaling to the margins of these colonies, combined with the presumptive role of BMP inhibitors secreted by the colony interior and consistent with a reaction-diffusion model (Etoc et al., 2016; Tewary et al., 2017). By contrast, hESCs plated at high densities (3,000-4,000 cells/mm^2^) using “funnels” on laminin-rich reconstituted basement membrane (rBM) modified polyacrylamide hydrogels (PA; E = 10^2^ to 10^5^ Pa) self-assemble disc-shaped colonies of hESCs and exhibit dramatically enhanced mesoderm specification upon BMP4 stimulation when cultured on the soft (E = 10^2^ to 10^3^ Pa) versus the stiff (E = 10^4^ to 10^5^ Pa) substrates (L. Przybyla et al., 2016; Laralynne Przybyla et al., 2016). An elastic modulus of 10^2^ to 10^3^ Pa is of the order of magnitude estimated for soft tissues, such as would be anticipated for the early embryo. Indeed, atomic force microscopy (AFM) measurements of the gastrulation-stage chicken epiblast (Mikawa et al., 2004; Shahbazi et al., 2016), which resembles these hESC embryonic disc cultures in size and shape, yielded elasticity values within this range (Figure S1). Thus, to more directly investigate the role of ECM mechanics in the self-organized patterning of mesoderm specification in hESCs, we developed a system for BMP4-induced differentiation on embryo-like compliant PA gels (Figure 1A). For technical reasons (hydrogel tearing and z-displacements in E ∼10^2^ Pa; L. Przybyla et al., 2016) we chose the softest experimentally compatible PA gel (E = 2,700Pa).

**Figure 1:**
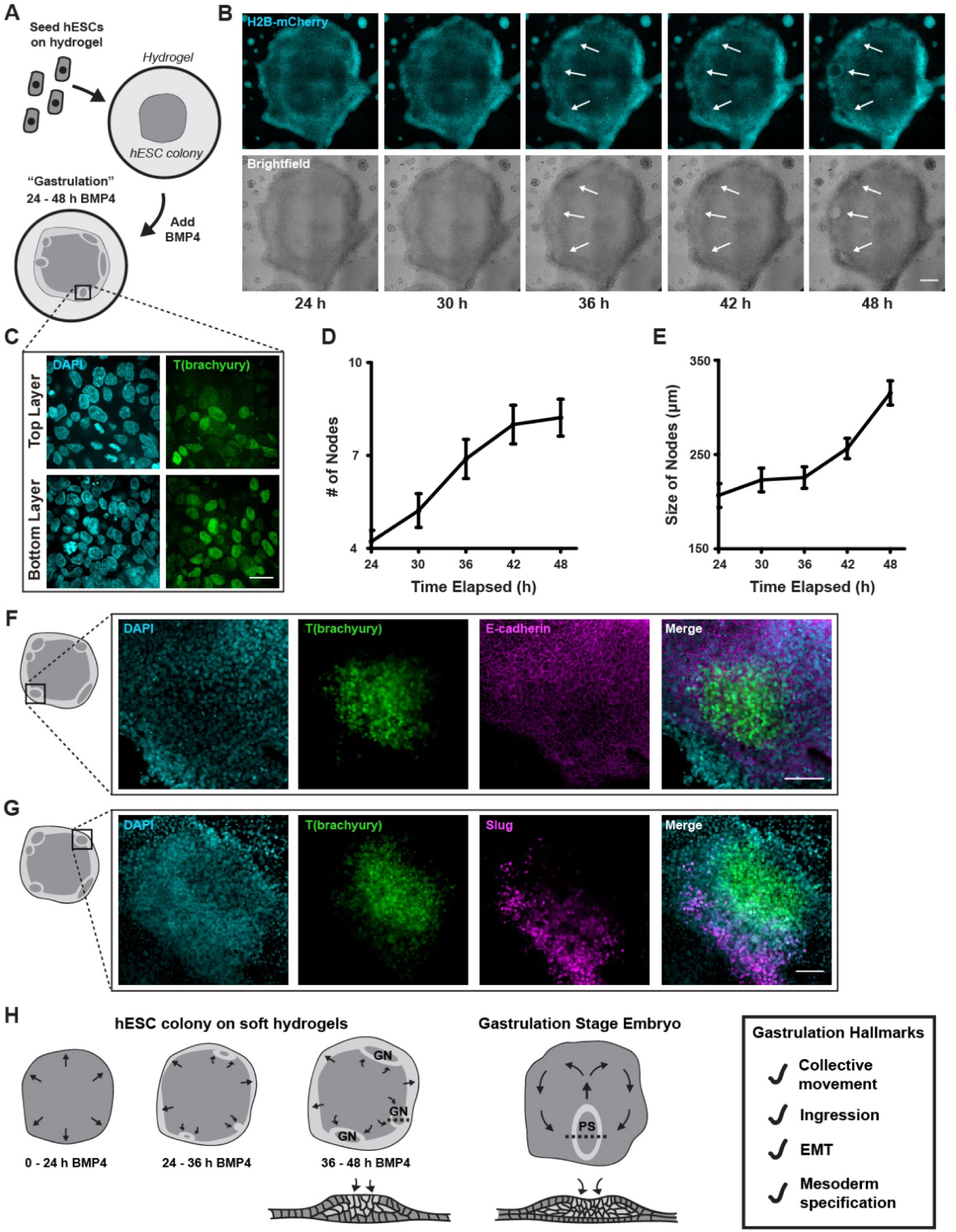
Compliant substrates promote hESC self-organization into “gastrulation-like” nodes. **A)** Cartoon of hESCs seeded as large colonies on compliant (2700 Pa) polyacrylamide hydrogels and stimulated with BMP4. **B)** Time-lapse brightfield (bottom) and immunofluorescent (top) images of an hESC colony stimulated with BMP4. Nuclei were visualized using H2B-mCherry (cyan). Arrows indicate “gastrulation-like” nodes. Scale bar = 500 μm. **C)** Representative spinning-disk confocal images of the top and bottom of the hESC colony 48 h following BMP4 stimulation showing T(brachyury) expression (mNeonGreen; green; right) and nuclei (DAPI; cyan; left) within the “gastrulation-like” nodes. The rectangle shown on the colony cartoon in panel A indicates the region of the hESC colony where the images were taken. Scale bar = 20 μm. **D)** Line graph quantifying the number of “gastrulation-like” nodes formed per hESC colony between 24 and 48 h after BMP4 stimulation. Each data point represents the mean number of nodes per colony ± SEM for n = 9 colonies from three independent experiments. **E)** Line graph quantifying the size of the “gastrulation-like” nodes formed between 24 and 48 h following BMP4 stimulation. Each data point represents the mean node size ± SEM of all the nodes identified in n = 9 colonies from three independent experiments. **F)** Representative immunofluorescent images of nuclei (DAPI; cyan; far left), T (mNeonGreen; green; middle-left), E-cadherin (Alexa568; magenta; middle-right), and composite (merge; far right) in the “gastrulation-like” nodes 48 h after BMP4 stimulation. The rectangle on the colony cartoon indicates the region within the colony where the images were taken. Scale bar = 100 μm. **G)** Representative immunofluorescent images of nuclei (DAPI; cyan; far left), T (mNeonGreen; green; middle-left), Slug (Alexa568; magenta; middle-right), and composite (merge; far right) in the “gastrulation-like” nodes 48 h after BMP4 stimulation. The rectangle on the colony cartoon indicates the region within the colony where the images were taken. Scale bar = 100 μm. **H)** Schematic representation of the “gastrulation-like” phenotype observed in the BMP4-stimulated hESC colonies plated on compliant hydrogels, compared to gastrulation in the developing embryo. Cross-sections along the dashed lines are depicted below the hESC colony and the developing embryo schematics. GN = gastrulation node. PS = primitive streak. EMT = epithelial to mesenchymal transition.

BMP4-induced differentiation in confined circular hESC colonies on ECM-patterned glass substrates specifies mesoderm within a concentric ring near the colony radial margins (Warmflash et al., 2014). Accordingly, we focused on the behavior of cells in the margins of unconfined circular colonies on the soft gels (E = 2,700 Pa). Live-imaging of H2B-mCherry-labelled nuclei following BMP4-mediated differentiation (50 ng/ml; 24 to 48 hours) revealed that discrete regions of highly dense hESCs formed near the periphery of the colonies (Figures 1B and 1D). High-magnification spinning disc confocal imaging showed that the hESCs collectively migrated to these nodes and further revealed that many ingressed basally upon reaching the node to ultimately assemble into a second cellular layer that expressed the mesoderm marker T(brachyury) (Figure 1C). This cellular behavior is highly reminiscent of gastrulation, wherein cells of the developing epiblast collectively migrate towards the midline and ingress to form a secondary transient structure called the primitive streak, which gives rise to the mesoderm and endoderm germ layers (Shahbazi et al., 2019; Shahbazi and Zernicka-Goetz, 2018; Simunovic and Brivanlou, 2017; Voiculescu et al., 2014; Williams and Solnica-Krezel, 2017). Analogous to the embryonic primitive streak that expands as gastrulation progresses, these “gastrulation-like” nodes in the hESC colonies similarly continued to expand between 24 and 48 hours post-BMP4-stimulation (Figure 1E). Moreover, and consistent with gastrulation, the hESCs that expressed T(brachyury) in “gastrulation-like” nodes also lost E-cadherin, indicating they had undergone an epithelial to mesenchymal transition (EMT; Figure 1F). Indeed, the radially-migrating cells adjacent to the “gastrulation-like” nodes expressed another EMT marker, Slug, implying they were in the process of ingression and undergoing mesoderm specification (Figure 1G).

These findings illustrate our ability to reproducibly induce a “gastrulation-like” phenotype, indicative of early embryogenesis, in self-organized nodes of hESCs that bear a striking similarity to the primitive streak that forms during gastrulation in the embryo (Figure 1H). The radial organization of mesoderm specification that we observed in our hESC model bears some similarity to the radial organization documented in the earlier studies of hESCs plated on patterned glass substrates (Etoc et al., 2016; Tewary et al., 2017; Warmflash et al., 2014). However, the formation of discrete “gastrulation-like” nodes in unconfined hESC colonies on PA gels is distinct from the apparent continuous concentric ring patterns of the primary germ layers observed in these prior models, and instead, is morphologically more reminiscent of the discrete primitive streak. These distinctions likely reflect differences in the biophysical properties of the local microenvironment, as well as constraints on outward radial cell migration imposed on confined (non-permissive) versus unconfined (permissive) colonies. The findings underscore the importance of the biophysical properties of the local microenvironment in recapitulating self-organization of the early embryo.

### Real-time monitoring of hESC “gastrulation-like” nodes

To explore the mechanisms regulating the self-organization that fosters the “gastrulation-like” phenotype in hESCs cultured on compliant substrates, we built a T-reporter hESC line using CRISPR homology-directed (HDR) repair (Chu et al., 2015; San Filippo et al., 2008) to facilitate monitoring the temporal development of the nodes in real-time (Figure 2A). After confirming that only the hESCs expressing T-mNeonGreen stained positive for the endogenous T protein following BMP4 differentiation (Figure 2B and 2C), we conducted gene expression analysis on fluorescence-activated cell sorted (FACS) populations of the T-reporter hESCs post-BMP4-stimulation (36 hours) to assess the fate commitment of the gastrulating cells (Figure 2D). Quantitative polymerase chain reaction (qPCR) revealed that the isolated T-positive hESCs expressed high levels of the mesoderm markers T(brachyury) and goosecoid (GSC), and the EMT marker Snai2, as compared to T-negative hESCs (Figure 2E). Additionally, we found that direct transcriptional targets of T (Koch et al., 2017) were upregulated in the T-positive hESCs, verifying that the T-mNeonGreen fusion protein did not compromise downstream transcriptional activity of T (Figure S2). By contrast, T-negative hESCs expressed high levels of the pluripotent marker Sox2, which also signifies ectodermal fate following gastrulation (Figure 2E; Koch et al., 2017; Warmflash et al., 2014). These data confirm that mesoderm specification occurred exclusively in the observed “gastrulation-like” nodes, and indicate that the cells ingressing within these nodes undergo transcriptional changes akin to the cells that pass through the primitive streak during early gastrulation.

**Figure 2:**
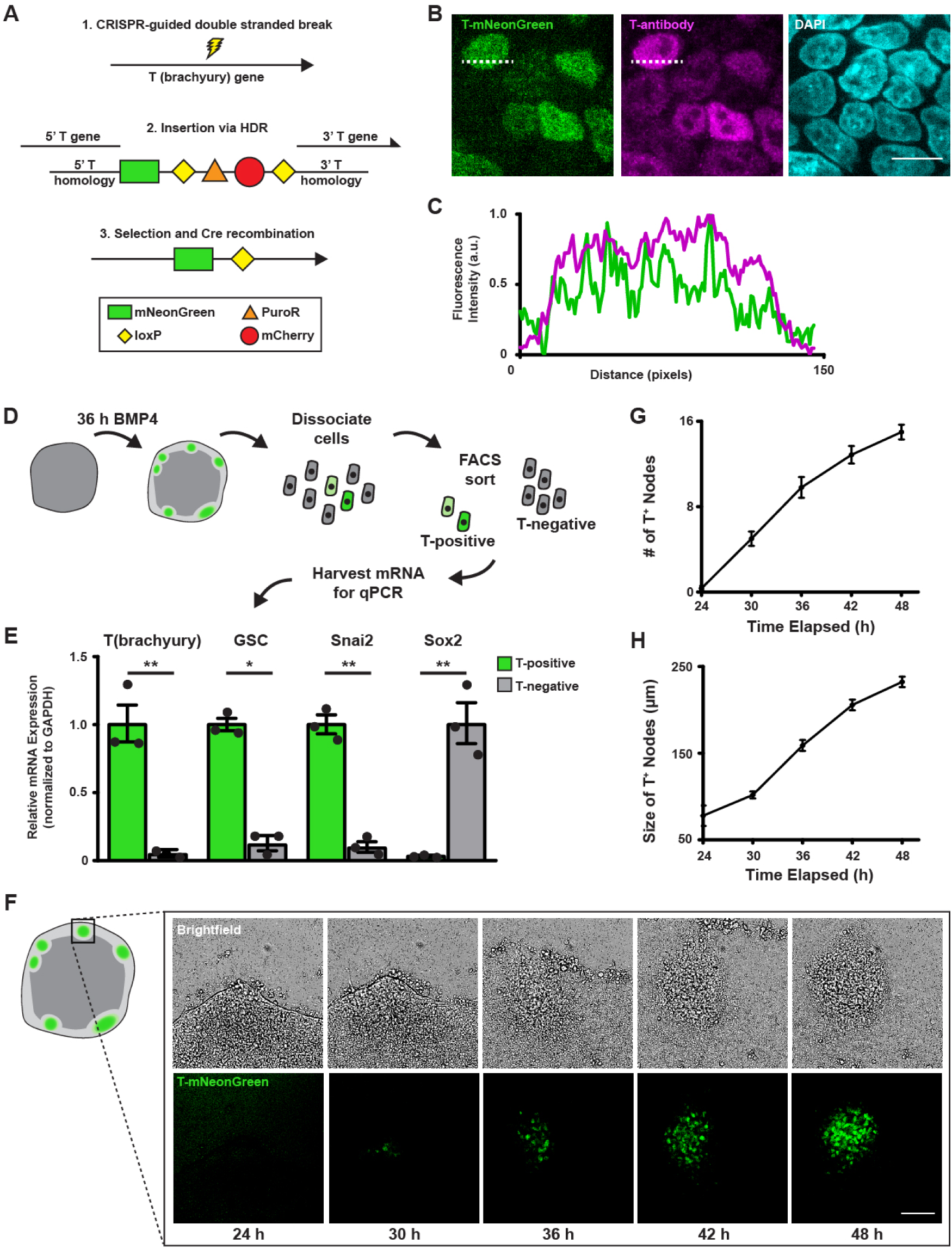
Real-time monitoring of hESC “gastrulation-like” nodes. **A)** Schematic representation of the T-mNeonGreen reporter system built using CRISPR-HDR. **B)** Representative immunofluorescent images of T(brachyury) labeled by the T-reporter (mNeonGreen; green; left) and an antibody to T (Alexa568; magenta; middle), and nuclei (DAPI; cyan; right). **C)** Plot of the fluorescence intensity of T-mNeonGreen (green line) and antibody-labeled T (magenta line) measured along the dashed lines shown in panel B. Scale bar = 10 μm. **D)** Cartoon showing the protocol used to compare gene expression between T-mNeonGreen-positive and T-mNeonGreen-negative cells 36 h following BMP4 stimulation. **E)** Bar graphs showing mRNA levels of mesoderm genes in T-positive and T-negative cells isolated from hESC colonies 36 h following BMP4 stimulation. Bars represent mean fold change ± SEM from three independent experiments. *p < 0.05 and **p < 0.01. **F)** Representative time-lapse brightfield (top) and immunofluorescent (T-mNeonGreen; green; bottom) images of a “gastrulation-like” node between 24 and 48 h after BMP4 stimulation. The rectangle on the colony cartoon indicates the region within the hESC colony where the images were taken. Scale bar = 100 μm. **G)** Line graph quantifying the number of “gastrulation-like” nodes marked by T-mNeonGreen per hESC colony between 24 and 48 h after BMP4 stimulation. Each data point represents the mean number of nodes per colony ± SEM for n = 15 colonies from three independent experiments. **H)** Line graph quantifying the size of the “gastrulation-like” nodes marked by T-mNeonGreen between 24 and 48 h after BMP4 stimulation. Each data point represents the mean node size ± SEM of all the nodes identified in n = 15 colonies from three independent experiments. HDR = homology-directed repair. a.u. = arbitrary units. GSC = goosecoid.

Importantly, time-course imaging of T-reporter hESCs following BMP4 stimulation revealed that the dramatic dynamic behavior of the entire tissue-like structure fosters the induction and expansion of the “gastrulation-like” nodes (Figure 2F). Coincident with T-expression and emergence of the “gastrulation-like” phenotype, the hESCs approximately 50 microns or 10 cell diameters inward from the colony periphery begin to assemble into densely packed aggregates (Figures 1F, 2F, and 2G) reminiscent of cells with enhanced cell-cell adhesions (Lakins et al., 2012; Laralynne Przybyla et al., 2016). These densely packed cellular aggregates progressively increased in size (Figure 2H), presumably due to observed coordinated migration of a subpopulation of cells initially moving radially outwards from the colony periphery that circle back inwards to contribute to the developing “gastrulation-like” nodes (Figure 2F). These collective movements bear a striking resemblance to the “Polonaise” movement that drives epiblast cells toward the midline and establishes the primitive streak in the gastrulating chick embryo (Voiculescu et al., 2014). These data thus provide additional compelling evidence that recapitulation of the compliance of the embryonic microenvironment enables self-organization of hESCs to facilitate coordinated programs of early embryogenesis and tissue development (Figure 1H).

Consistent with this paradigm, live-cell imaging also identified a second population of cells with a distinct migration and adhesion phenotype. This second cell population migrated outward from the colony periphery and exhibited a flattened morphology suggestive of high cell-ECM adhesion activity (Figure 2F). The distinct morphologies displayed by these two hESC populations is reminiscent of the differential cell-cell and cell-ECM adhesion phenotype that regulates branching morphogenesis, drives epithelial cyst morphogenesis, and permits efficient mesoderm specification in hESCs (Affolter et al., 2003; Nelson et al., 2006; Paszek et al., 2005; Paszek and Weaver, 2004; Laralynne Przybyla et al., 2016; Weaver et al., 2002). The findings imply that the differential cell-cell versus cell-ECM tension fostered by growing the hESCs on compliant hydrogels contributes to tissue self-organization and facilitates the “gastrulation-like” phenotype.

### Cell-cell tension directs “gastrulation-like” node organization to specify mesoderm

We next asked whether a causal link exists between cell-cell tension fostered in hESC colonies on compliant substrates and the “gastrulation-like” tissue self-organization. We measured cell-adhesion tension using traction force microscopy (TFM) and registered these forces with the emerging “gastrulation-like” nodes using live-cell imaging of the T-reporter (L. Przybyla et al., 2016). Remarkably, the BMP4-stimulated “gastrulation-like” nodes developed within the same hESC colony regions that exhibited the highest cell-adhesion tension (Figures 3A and 3B). Moreover, and importantly, the elevated cell-adhesion tension arose prior to mesoderm specification, as indicated by the T-reporter (Figure 2F). The data thus imply that cell-adhesion-dependent tissue-level force regulates tissue self-organization to spatially direct the “gastrulation-like” phenotype.

**Figure 3:**
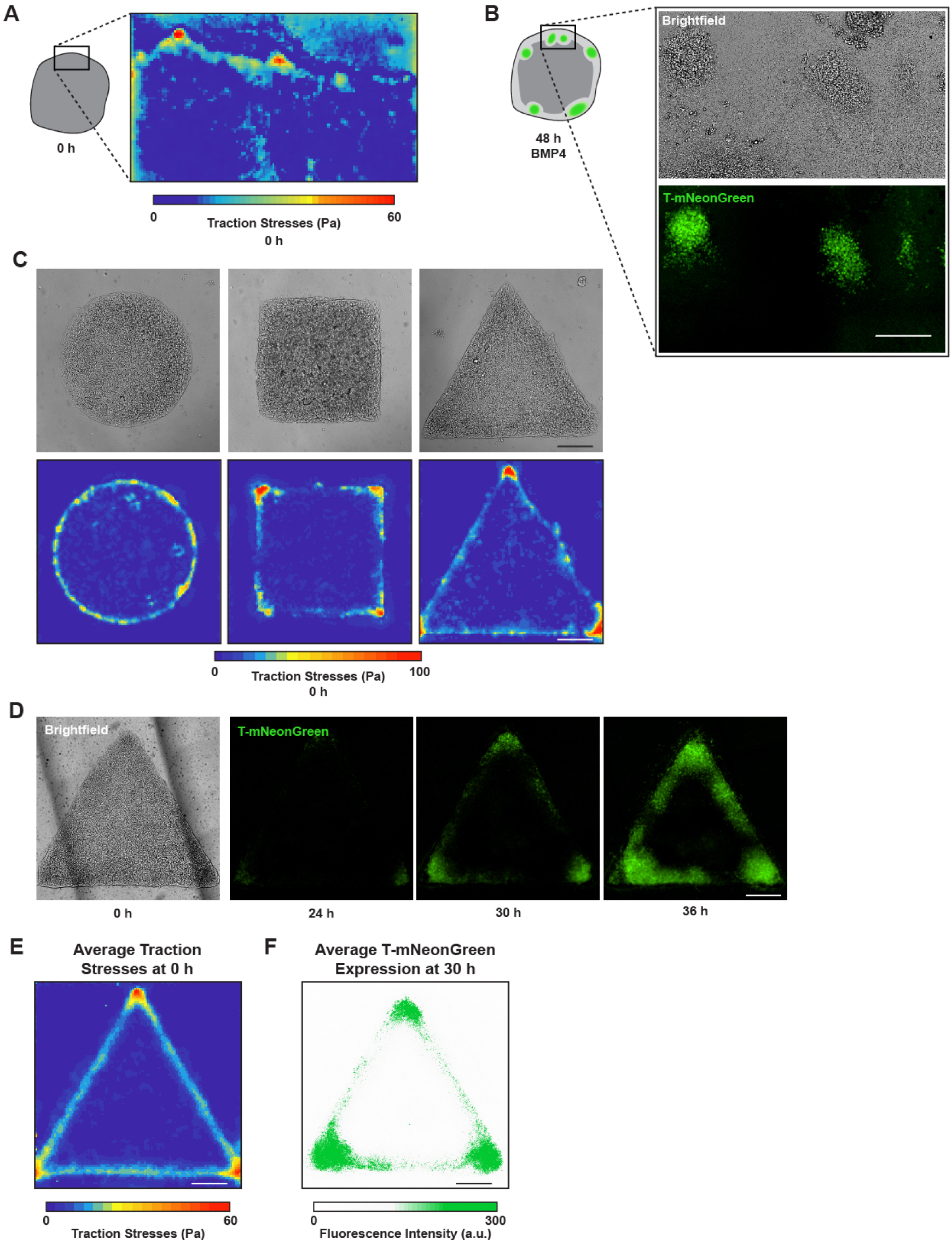
Cell-cell tension directs “gastrulation-like” node organization to specify mesoderm. **A)** Map of traction stresses generated at the periphery of a representative unconfined hESC colony prior to BMP4 stimulation. The rectangle on the colony cartoon indicates the region within the hESC colony where the traction stresses were measured. **B)** Brightfield (top) and immunofluorescent (T-mNeonGreen; green; bottom) images 48 h after BMP4 stimulation in the same field of view where the traction stresses were measured in panel A. **C)** Representative brightfield images (top) and corresponding traction stress maps (bottom) measured in the geometrically-confined hESC colonies plated on the compliant gels before BMP4 stimulation (0 h). **D)** Representative brightfield image and time-lapse T-mNeonGreen immunofluorescent images of a geometrically-confined triangle hESC colony before (0 h; brightfield) and after BMP4 stimulation (24 h, 30 h, 36 h; T-mNeonGreen). **E)** Map of average traction stresses measured for the geometrically-confined triangle hESC colonies. n = 19 colonies from three independent experiments. **F)** Cumulative intensity map of average T-mNeonGreen expression within the geometrically-confined triangle hESC colonies 30 h after BMP4 stimulation. n = 12 colonies from four independent experiments. All scale bars = 250 μm. Pa = Pascals. a.u. = arbitrary units.

To test the relationship between tissue-level force and gastrulation, we patterned the geometry of hESC colonies on compliant substrates designed to engineer the localization of cell-adhesion tension (Muncie et al., 2019). After confirming that hESCs plated on the patterned substrates did not spontaneously differentiate (Figure S3), we monitored the force-generating behavior of the hESC colonies. We determined that specific tissue geometries previously shown to promote localized tension within an epithelial cell colony (Gomez et al., 2010; Kilian et al., 2010; Lee et al., 2016; Nelson et al., 2005; Smith et al., 2018) similarly induced high tension in the hESC colonies. For instance, shapes such as squares and triangles fostered the highest cell tension development in the colony corners, whereas circles developed comparatively moderate levels of tension around their colony periphery (Figure 3C). As predicted, live-cell imaging of the T-reporter showed that following BMP4 stimulation, “gastrulation-like” nodes arose in the colony regions that displayed the highest cell-adhesion tension (Figure 3D). Moreover, the averaged intensity of multiple cell tension and T-expression plots, generated through cumulative time-lapse imaging of T-reporter activity together with cell-adhesion tension measurements across multiple colonies, confirmed that mesoderm specification conferred by the “gastrulation-like” nodes is indeed spatially patterned by regions of high cell-adhesion tension (Figures 3E and 3F). Accordingly, the data reveal that regions of high cell-adhesion tension that develop within a tissue-like structure direct the spatial localization of “gastrulation-like” nodes that, in turn, drive mesoderm specification.

### Cell-cell adhesion mediates the high cell tension required to specify “gastrulation-like” nodes

To further explore the role of cell-adhesion-mediated tissue-level tension in hESC colony self-organization and “gastrulation-like” node formation, we attenuated cell-cell adhesions and examined the effect on mesoderm specification. Inducible short hairpin RNA knockdown that achieved a 50 percent reduction in E-cadherin (shE-cadherin; Figures 4A to 4C) was sufficient to significantly reduce the magnitude of cell-adhesion tension in the triangle patterned hESC colonies relative to control colonies, which was particularly evident in the colony corners (Figures 4D and 4E). This level of E-cadherin knockdown and the reduced tissue tension also significantly decreased the level of mesoderm specification observed in the hESC colonies (Figures 4F and 4G). The findings indicate that cell-cell adhesion, mediated by molecules such as E-cadherin, is necessary for the generation of spatially restricted regions of high colony cell-adhesion tension that in turn directs mesoderm specification at the “gastrulation-like” nodes in the hESC colonies.

**Figure 4:**
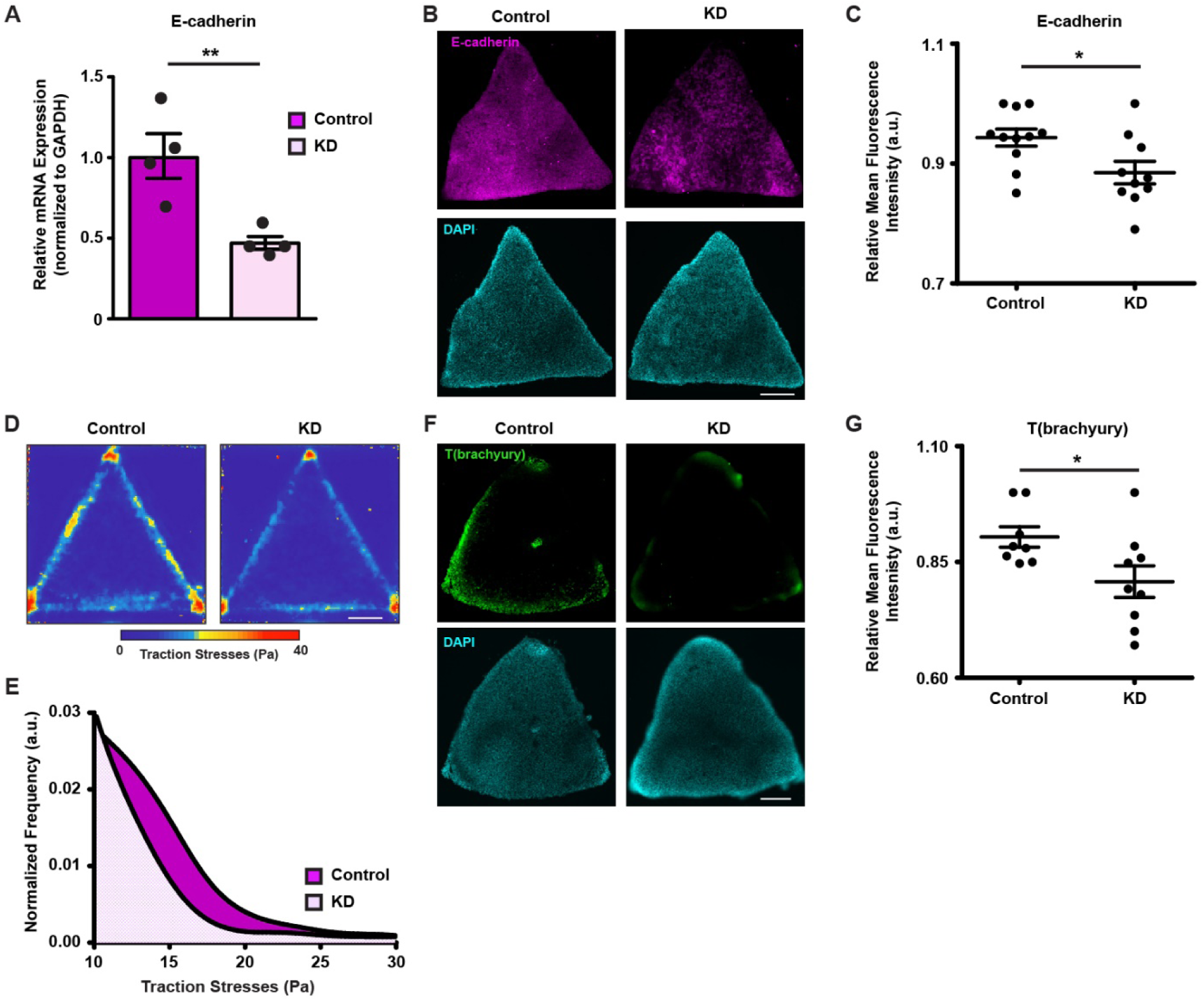
Cell-cell adhesion mediates the high cell tension required to specify “gastrulation-like” nodes. **A)** Bar graph showing mRNA expression of E-cadherin in the geometrically-confined colonies of hESCs with (KD) and without (control) shE-cadherin knockdown. Bars represent mean fold change ± SEM from four independent experiments. **p < 0.01. **B)** Representative immunofluorescent images of E-cadherin (Alexa568; magenta; top) and nuclei (DAPI; cyan; bottom) in geometrically-confined triangle hESC colonies with (KD; right) and without (control; left) shE-cadherin knockdown. **C)** Scatter plot quantifying immunofluorescent signal of E-cadherin in images represented by panel B. Individual points represent fluorescence intensity measured for a single image. Bars represent relative fluorescence intensity mean ± SEM for images taken from three independent experiments. *p < 0.05. **D)** Map of average traction stresses measured within geometrically-confined triangle hESC colonies with (KD; right) and without (control; left) shE-cadherin knockdown. n = 13 colonies for each condition from three independent experiments. **E)** Kernel distribution plot of average traction stresses for the geometrically-confined triangle hESC colonies with (KD) and without (control) shE-cadherin knockdown. **F)** Representative immunofluorescent images of T (Alexa488; green; top) and nuclei (DAPI; cyan; bottom) 36 h after BMP4 stimulation in geometrically-confined hESC triangle colonies with (KD; right) and without (control; left) shE-cadherin knockdown. **G)** Scatter plot quantifying immunofluorescent signal of T(brachyury) in images represented by panel F. Individual points represent fluorescence intensity measured for a single image. Bars represent relative fluorescence intensity mean ± SEM for images taken from three independent experiments. *p < 0.05. All scale bars = 250 μm. KD = knockdown. Pa = Pascals. a.u. = arbitrary units.

### Ablating regions of high cell tension inhibits the development of “gastrulation-like” nodes

To definitively establish a role for high cell-cell and tissue-level tension in colony self-organization and induction of the “gastrulation-like” behavior, we directly ablated tension in patterned colonies of hESCs. We generated precise cuts across the corners of the triangle hESC colonies using an eyebrow knife (Sive et al., 2000), which TFM revealed disrupted the spatially-restricted regions of high cell-adhesion tension (Figures 5A and 5B). Consistent with our assertion that localized regions of high cell-adhesion tension direct the “gastrulation-like” nodes that promote mesoderm specification, we noted a significant delay in BMP4-induced T-reporter expression in the cut corners of these colonies (Figures 5C and 5D). To provide additional evidence of a causal link between localized cell-adhesion tension and development of the “gastrulation-like” nodes, we next constructed Pac-Man patterned surfaces that generate low cell tension at the concave edge of the hESC colony, corresponding to the “mouth” of the Pac-Man (Figures 5E and 5F). Critically, although BMP4-stimulation of these patterned Pac-Man hESC colonies induced T-reporter expression at the convex edges of the Pac-Man, T-reporter expression was clearly excluded from the low-tension concave edge (Figures 5E to 5G). The results provide compelling evidence that high cell-adhesion tension directs the spatially restricted development of “gastrulation-like” nodes that specify mesoderm in these hESC colonies.

**Figure 5:**
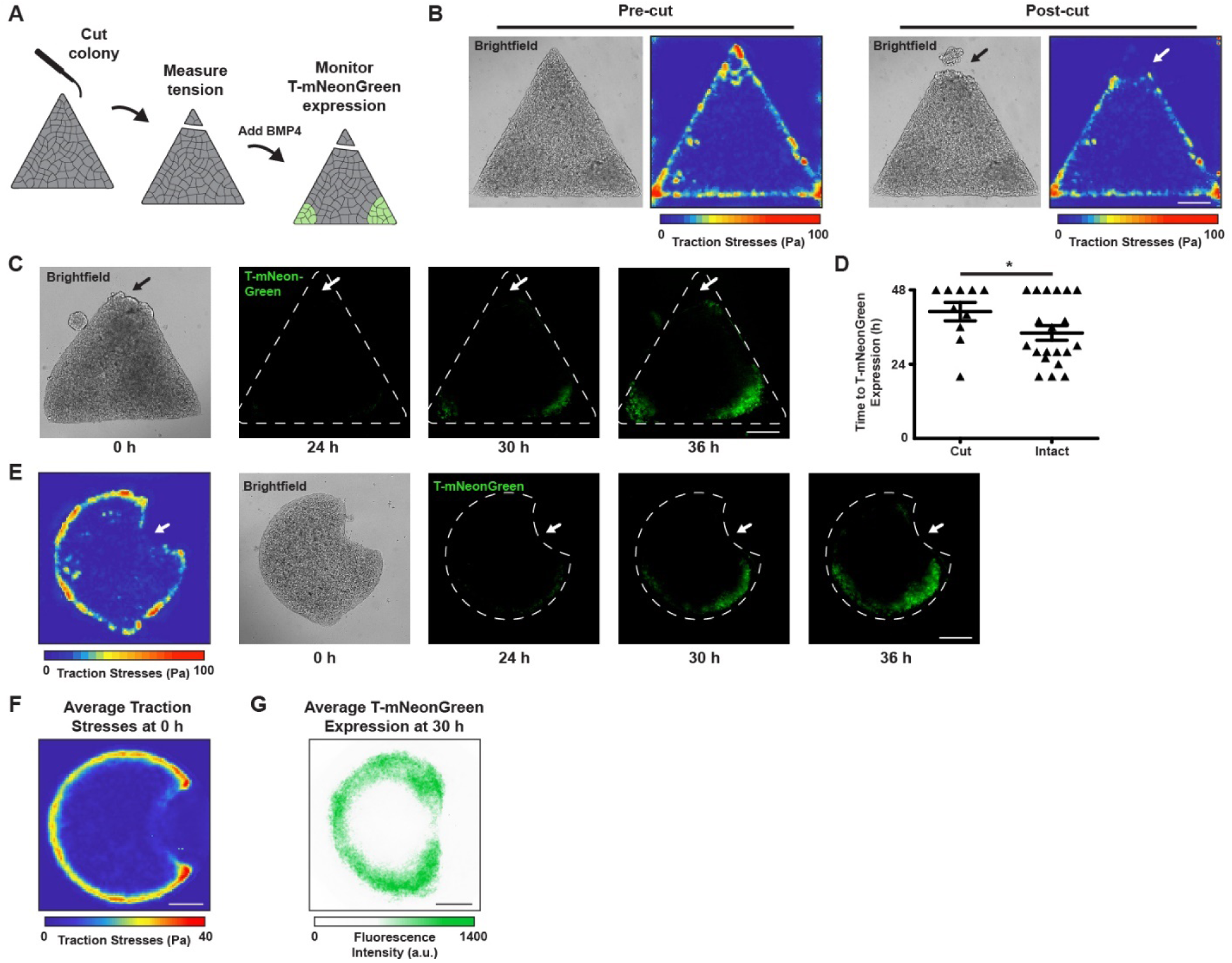
Ablating regions of high cell tension inhibits the development of “gastrulation-like” nodes. **A)** Cartoon of eyebrow knife experiment. **B)** Representative brightfield images (left panels) and traction stress maps (right panels) of the geometrically-confined triangle hESC colonies before and after eyebrow knife ablation across a colony corner. **C)** Representative brightfield image and time-lapse T-mNeonGreen immunofluorescent images of a geometrically-confined triangle hESC colony whose corner has been ablated using the eyebrow knife, before (0 h; brightfield) and after BMP4 stimulation (24 h, 30 h, 36 h; T-mNeonGreen). Arrow indicates the site of eyebrow knife ablation. **D)** Scatter plot quantifying immunofluorescently detected T-mNeonGreen in the geometrically-confined triangle hESC colonies with and without eyebrow knife ablation between 0 and 48 h following BMP4 stimulation. Bars represent mean ± SEM for data from n = 10 colonies from three independent experiments. *p < 0.05. **E)** Representative traction stress map and corresponding brightfield and time-lapse T-mNeonGreen immunofluorescent images in a typical geometrically-confined Pac-Man hESC colony before (0 h; traction stress map and brightfield) and after (24 h, 30 h, 36 h; T-mNeonGreen) BMP4 stimulation. Arrow indicates concave edge, “mouth” of Pac-Man. **F)** Map of average traction stresses measured within geometrically-confined Pac-Man hESC colonies. n = 20 colonies from four independent experiments. **G)** Cumulative intensity map of average T-mNeonGreen expression within the geometrically-confined Pac-Man hESC colonies 30 h after BMP4 stimulation. n = 20 colonies from four independent experiments. All scale bars = 250 μm. Pa = Pascals. a.u. = arbitrary units.

### High cell tension promotes β-catenin release from adherens junctions to specify mesoderm

Mesoderm specification in hESC colonies depends on Wnt/β-catenin signaling and is enhanced by growth on a compliant matrix (Laralynne Przybyla et al., 2016). Accordingly, we examined whether localized cell-adhesion tension specifies mesoderm by promoting the release of β-catenin from E-cadherin adhesion complexes, specifically in regions of high tension. We observed the preferential loss of β-catenin from E-cadherin junctions within the localized regions of high cell-adhesion tension 24 hours following BMP4 stimulation (Figure 6A). Activated phospho-Src-family kinases (pSFKs) phosphorylate junctional β-catenin to facilitate its release from adherens junction complexes (Bienz, 2005; Gayrard et al., 2018; Gottardi and Gumbiner, 2004; Howard et al., 2011; Lilien and Balsamo, 2005; Laralynne Przybyla et al., 2016). Consistently, we detected high levels of pSFKs within 6 hours of BMP4 stimulation, specifically within regions of high cell-adhesion tension (Figure 6B). Furthermore, blocking pSFK activity, using the inhibitor PP1, prevented β-catenin release at the E-cadherin junctions within the regions of high tension (Figures 6B and 6C). Preventing β-catenin release resulted in a significant loss of mesoderm specification, as indicated by decreased T(brachyury), goosecoid, and Snai2 expression, and elevated Sox2 levels (Figure 6D).

**Figure 6:**
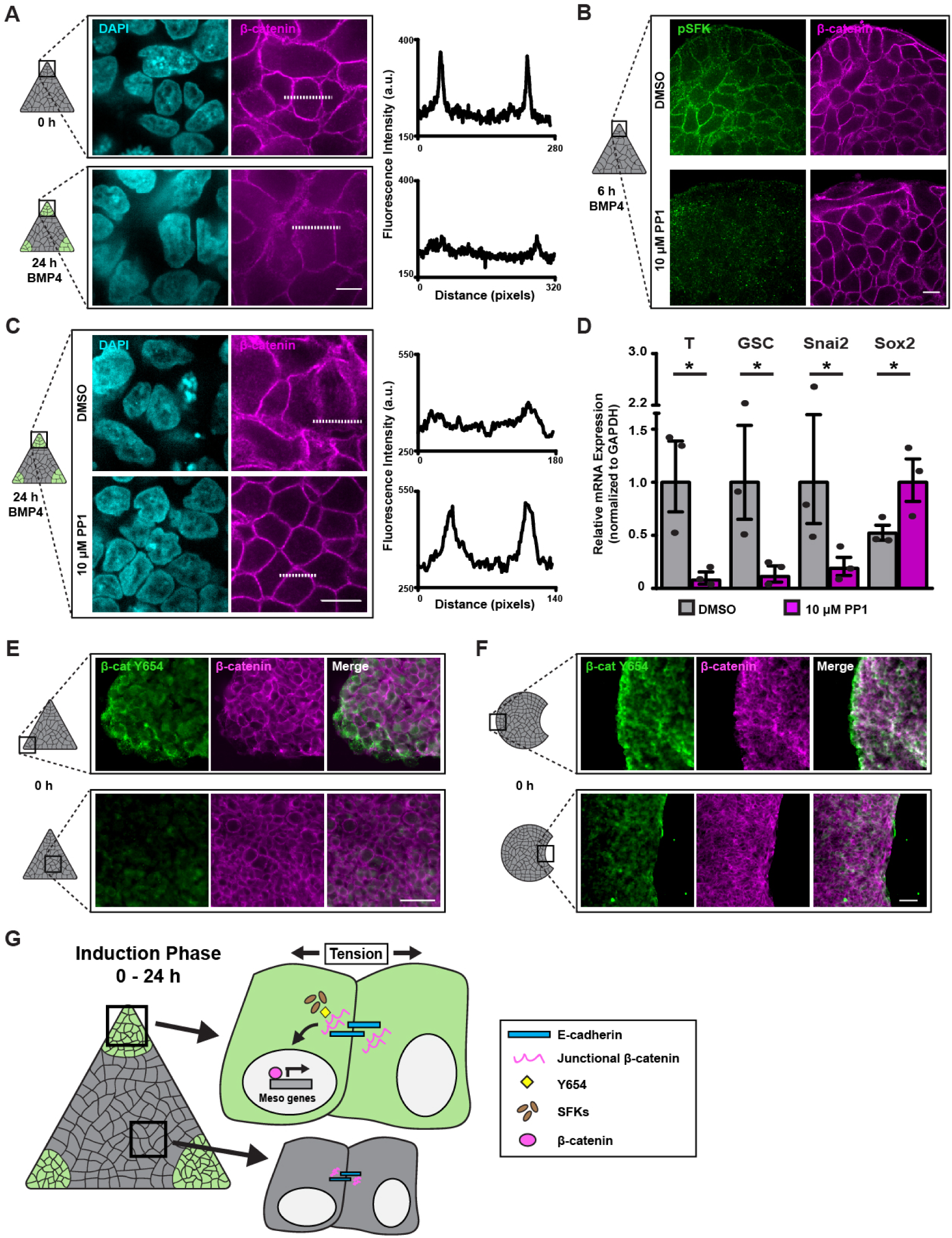
High cell tension promotes β-catenin release from adherens junctions to specify mesoderm. **A)** Representative immunofluorescent images of nuclei (DAPI; cyan; left) β-catenin (Alexa568; magenta; right) in the corners of a typical geometrically-confined triangle hESC colony before (top) and 24 h after (bottom) BMP4 stimulation. The intensity of β-catenin fluorescence was measured along the dashed lines and plotted (far right). Scale bar = 10 μm. **B)** Representative immunofluorescent images of phosphorylated Src-family kinases (Alex488; green; left) and β-catenin (Alexa568; magenta; right) in the corners of a typical geometrically-confined triangle hESC colony 6 h after stimulation with BMP4 and treatment with vehicle (DMSO; top) or Src inhibitor (PP1; 10 μM; bottom). Scale bar = 10 μm. **C)** Representative immunofluorescent images of nuclei (DAPI; cyan; left) and β-catenin (Alexa 568; magenta; right) in the corners of a typical geometrically-confined triangle hESC colony 24 h after BMP4 stimulation and treatment with vehicle (DMSO; top) or Src inhibitor (PP1; 10 μM; bottom). The intensity of β-catenin fluorescence was measured along the dashed lines and plotted (far right). Scale bar = 10 μm. **D)** Bar graphs showing mRNA levels of mesoderm genes expressed in the geometrically-confined hESC colonies 36 h after BMP4 stimulation and treatment with either vehicle (DMSO) or Src inhibitor (PP1; 10 μM). Bars represent mean fold change ± SEM from three independent experiments. *p < 0.05. **E)** Representative immunofluorescent images of β-catenin Y654 (Alex488; green; left), total β-catenin (Alexa 568; magenta; middle), and composite (merge; right) in a typical geometrically-confined triangle hESC colony prior to BMP4 stimulation. Scale bar = 50 μm. **F)** Representative immunofluorescent images of β-catenin Y654 (Alex488; green; left), total β-catenin (Alexa 568; magenta; middle), and composite (merge; right) in a typical geometrically-confined Pac-Man hESC colony prior to BMP4 stimulation. Scale bar = 50 μm. **G)** Cartoon summarizing the mechanism by which regionally-localized high cell-adhesion tension exposes β-catenin Y654 to facilitate its subsequent phosphorylation by pSFKs following BMP4 stimulation. Upon phosphorylation by pSFKs, β-catenin is released from the E-cadherin junctions and thereafter translocates to the nucleus where it regulates gene transcription to induce mesoderm specification in the regions of high tissue tension. The rectangles on the colony cartoons indicate the regions within the colonies where the images were taken. β-cat Y654 = tyrosine 654 of β-catenin. pSFK = phosphorylated Src-family kinases. a.u. = arbitrary units.

Molecular dynamic simulations showed tension across E-cadherin junctions exposes the tyrosine 654 (Y654) of cadherin-bound β-catenin (Röper et al., 2018). Therefore, we asked whether high cell-adhesion tension fosters a conformational change in β-catenin that permits its phosphorylation on Y654 to spatially pattern mesoderm specification at the nodes of high tension. Consistently, we detected enhanced binding of an antibody specific to the β-catenin Y654 in the cells at the corners of the triangle patterned hESC colonies, where we measured higher cell-adhesion tension, suggesting that tension exposed the Y654 phosphorylation site (Figure 6E). The increased tyrosine Y654 accessibility induced by the high cell-adhesion tension subsequently permitted SFK-mediated phosphorylation (Figure 6B) and release of β-catenin from the E-cadherin junctions (Figure 6A) to promote mesoderm specification. Furthermore, in the Pac-Man patterned hESC colonies, Y654 was similarly exposed at the convex edges of the colony, in the regions experiencing high cell-adhesion tension, whereas the concave edge experiencing low tension had low to non-detectable Y654 accessibility (Figure 6F). The data argue that high cell-adhesion tension specifies the spatial localization of “gastrulation-like” nodes that give rise to mesoderm by modifying the structure-function of a key transcriptional regulator, β-catenin, whose accessibility and nuclear translocation are regulated by SFK-mediated phosphorylation (Figure 6G).

### Wnt signaling reinforces mesoderm specification in regions of high tension

To verify that cell tension specifies mesoderm by regulating Wnt/β-catenin signaling, we used FACS to isolate T-positive cells from patterned hESC colonies following BMP4 stimulation (36 hours; subsequent to β-catenin release from the E-cadherin junctions), and looked for evidence of Wnt signaling (Figure 7A). Gene expression analysis revealed that the canonical Wnt ligands Wnt3a and Wnt8a, which are critical for mesoderm specification (Chhabra et al., 2019; Kemp et al., 2005; Lindsley et al., 2006; Martyn et al., 2019; Laralynne Przybyla et al., 2016), were expressed at significantly higher levels in the T-positive cells isolated from the regions of high cell-adheesion tension (Figure 7B). By contrast, levels of the non-canonical ligand Wnt4 were higher in the isolated T-negative cells (Figure 7B). The findings reveal that in response to BMP4, β-catenin induces mesoderm-promoting Wnts exclusively in regions of high cell tension.

**Figure 7:**
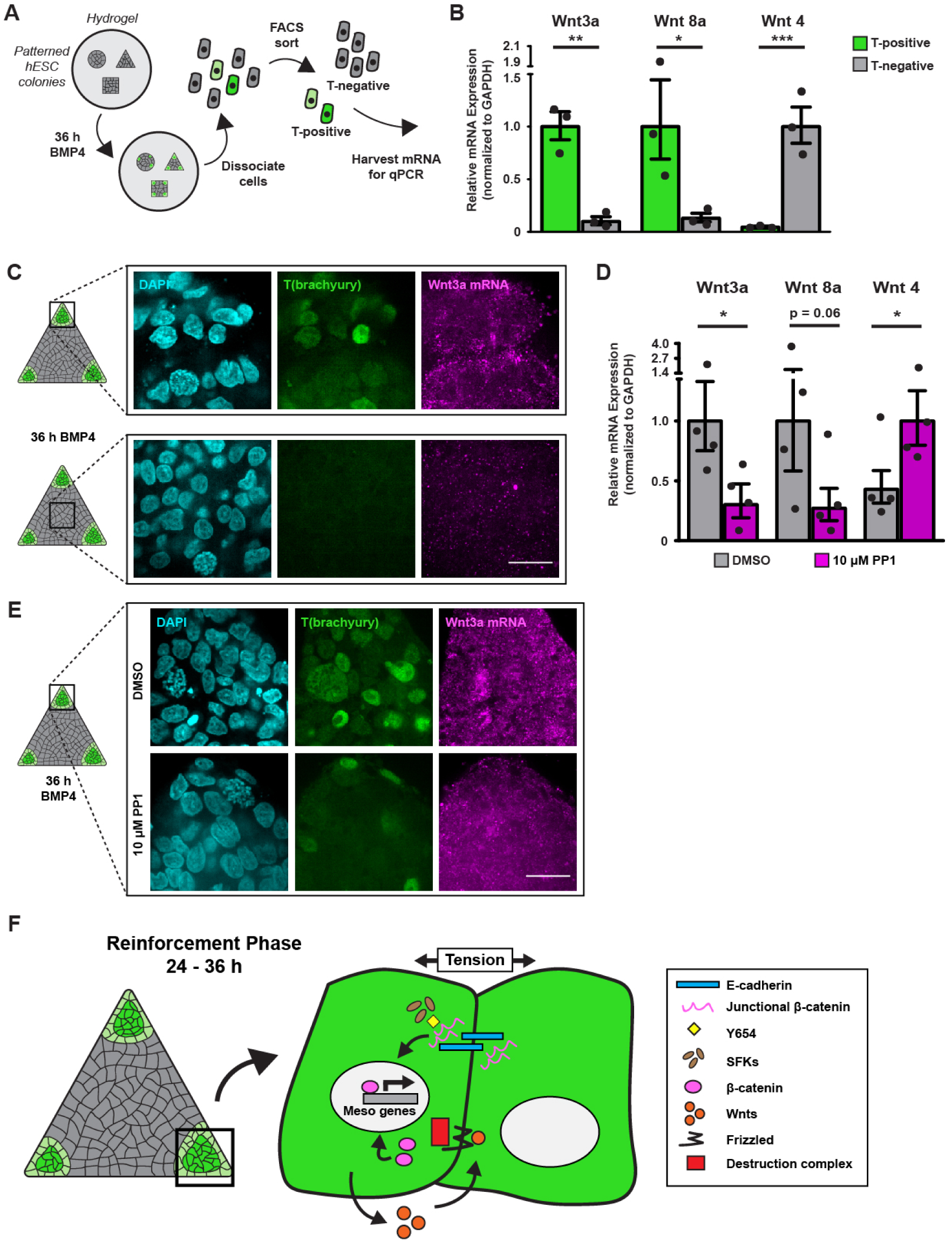
Wnt signaling reinforces mesoderm specification in regions of high tension. **A)** Cartoon of the protocol used to compare Wnt expression in the T-mNeonGreen-positive and T-mNeonGreen-negative hESCs 36 h after BMP4 stimulation. **B)** Bar graphs showing mRNA levels of Wnt ligands in the T-positive and T-negative hESCs isolated from the geometrically-confined hESC colonies 36 h after BMP4 stimulation. Bars represent mean fold change ± SEM from three independent experiments. *p < 0.05, **p < 0.01, and ***p < 0.001. **C)** Representative immunofluorescent images of nuclei (DAPI; cyan; left), immune-detected T(brachyury) protein (mNeonGreen; green; middle), and *in-situ*-detected Wnt3a mRNA (Alexa647; magenta; right) in regions corresponding to high (top) and low (bottom) cell-cell tension within geometrically-confined triangle hESC colonies 36 h after BMP4 stimulation. **D)** Bar graphs showing mRNA levels of Wnt ligands in the geometrically-confined hESC colonies 36 h after BMP4 stimulation and treatment with vehicle (DMSO) or Src inhibitor (PP1; 10 μM). Bars represent mean fold change ± SEM from four independent experiments. *p < 0.05. **E)** Representative immunofluorescent images of nuclei (DAPI; cyan; left), immune-detected T(brachyury) protein (mNeonGreen; green; middle), and *in-situ*-detected Wnt3a mRNA (Alexa647; magenta; right) in regions corresponding to high cell-cell tension within geometrically-confined triangle hESC colonies 36 h after BMP4 stimulation and treatment with vehicle (DMSO; top) or Src inhibitor (PP1; 10 μM; bottom). **F)** Cartoon summarizing the mechanism by which regions of high cell-cell tension direct mesoderm specification in response to BMP4 stimulation. Following the Src-mediated release of junctional β-catenin, nuclear β-catenin enhances the expression of Wnt ligands within the spatially-localized nodes of high tension, and these Wnt ligands thereafter promote mesoderm specification via a feed-forward circuit. The rectangles within the colony cartoons indicate the regions within the colonies where the fluorescent images were taken. All scale bars = 20 μm.

*In situ* hybridization chain reaction (ISH-HCR; Choi et al., 2018) verified that Wnt3a transcription was upregulated in response to BMP4 within the same high tension colony region, emphasizing a role for tissue-level force in mesoderm specification (Figures 7C and S4). Moreover, because blocking SFK activity with PP1 prevented the upregulation of the mesoderm-inducing Wnts, Src-mediated release and transcriptional activity of β-catenin appears to be necessary for tension-regulated Wnt ligand expression (Figures 7D and 7E). Collectively the findings elucidate a mechanism whereby high cell-adhesion tension promotes spatially-restricted gastrulation in hESC colonies by enhancing the expression of mesoderm-inducing Wnts through Src-mediated release of β-catenin from cadherin-catenin complexes (Figure 7F).

## Discussion

We demonstrated that high cell-adhesion tension directs the spatial localization of self-organizing “gastrulation-like” nodes in hESCs and does so by modulating Wnt/β-catenin signaling to induce transcription of mesoderm-specifying genes. It has long been recognized that mechanical forces are necessary to drive cell rearrangements and tissue organization within developing embryos (Hamada, 2015; Ko and Martin, 2020; Paré and Zallen, 2020; Voiculescu et al., 2014; Williams and Solnica-Krezel, 2017), and we provide compelling evidence that these same forces are capable of influencing the morphogen signaling pathways that regulate cell fate decisions by driving transcriptional reprogramming. This illustrates a critical interplay between the mechanics that drive tissue development and the molecular signaling that regulates cell fate decisions.

We identify a mechanism whereby regional high cell-adhesion tension modulates the conformation of β-catenin at cadherin-catenin complexes, permitting Src-mediated phosphorylation and release of β-catenin from cell junctions. The modified β-catenin is now free to translocate to the nucleus, where it exerts its transcriptional activity and induces the expression of multiple Wnt ligands that feed forward to promote and reinforce mesoderm specification and gastrulation (Bienz, 2005; Chhabra et al., 2019; Kemp et al., 2005; Lindsley et al., 2006; Martyn et al., 2019; Laralynne Przybyla et al., 2016). Notably, Wnt/β-catenin signaling is conserved among model organisms and plays multiple roles throughout the course of embryogenesis and in adult stem cells (Clevers, 2006; Eliazer et al., 2019; Petersen and Reddien, 2009; van Amerongen and Nusse, 2009), implying that the tension-regulated activation of the pathway that we illustrate here is likely also conserved and re-used throughout development. In fact, Farge and colleagues demonstrated that the same Src-mediated phosphorylation at Y654 and subsequent release of junctional β-catenin occurs during mesoderm invagination in *Drosophila* embryos (Röper et al., 2018). Moreover, a recent study of the tissue-level forces underlying gastrulation in avian embryos revealed that higher contractile forces are generated at the posterior of the embryo in the margin between the epiblast and extraembryonic tissue (Saadaoui et al., 2020). Interestingly, the geometric configuration of the gastrulating human embryo closely resembles that of these avian model organisms (Mikawa et al., 2004; Shahbazi et al., 2016). Given that we observe a strikingly similar tension-guided morphogenesis to avian gastrulation in hESC colonies, it seems likely that the tension-dependent modulation of the Wnt/β-catenin pathway we identified here may be a key regulator of both human and avian gastrulation.

Our work extends prior findings, which illustrated that variations in BMP receptor accessibility and secreted morphogen concentrations lead to patterning of mesoderm specification within a concentric ring inwards from the hESC colony edge (Chhabra et al., 2019; Etoc et al., 2016; Manfrin et al., 2019; Martyn et al., 2019; Tewary et al., 2019, 2017; Warmflash et al., 2014), to implicate force as an additional regulator of self-organization, and mesoderm specification in particular. Our studies expand upon these findings to suggest that colonies of hESCs cultured on compliant substrates are particularly useful for studying the physical and molecular regulators of early embryogenesis, specifically because they recapitulate the morphogenesis and transcriptional reprogramming that regulates gastrulation in the embryo. In contrast to a concentric ring of mesoderm specification, we observed self-organization of discrete “gastrulation-like” nodes that expand radially, distinctly resembling the formation and elongation of the primitive streak in the developing embryo (Mikawa et al., 2004; Voiculescu et al., 2014). Moreover, similar to epiblast cells that reach the primitive streak, our studies showed that the hESCs that contribute to these “gastrulation-like” nodes undergo EMT and ingress to form additional cell layers as they concomitantly begin expressing key mesoderm genes. The observed similarities between the “gastrulation-like” nodes in hESC colonies and the embryonic primitive streak suggest that this system could prove to be a powerful model with which to dissect the interplay between tissue force, cytoskeletal tension, and the signaling and transcriptional programs that regulate early development. Indeed, the cultured hESC system, combined with tissue patterning and tuned biophysical ECMs, lends itself to super-resolution imaging, single-cell sequencing techniques, quantitative force measurements, and physical manipulations that are challenging to perform using intact embryos, and which should yield critical fundamental insight into the process of human embryogenesis.

## Acknowledgements

We thank Lisandro Maya-Ramos and Takashi Mikawa for assistance with chicken embryo manipulation. We also thank Roberto Falcón-Banchs and Lydia L. Sohn for assistance with soft lithography techniques, which enabled the development of the method for geometric-confinement of hESCs on compliant substrates. Thanks to Dhruv Thakar for proofreading the manuscript. The authors would like to acknowledge funding from CIRM grant RB5-07409 and NIH/NCI U01 grant CA202241. Additionally, J.M.M. is thankful for support from the UCSF Discovery Fellowship.

## Author Contributions

Experimental Conceptualization, V.M.W., J.M.M., N.M.E.A., and J.N.L. Methodology, J.M.M. and J.N.L. Software, J.M.M. Validation, J.M.M. and J.N.L. Formal Analysis, J.M.M. Investigation, J.M.M., N.M.E.A., and J.N.L. Resources, V.M.W. Writing – Original Draft, J.M.M. and V.M.W. Writing – Review & Editing, V.M.W., J.M.M., N.M.E.A., and J.N.L. Visualization, J.M.M. Supervision, V.M.W. and J.N.L. Funding Acquisition, V.M.W.

## STAR Methods

### Lead Contact and Materials Availability

Further information and requests for resources and reagents should be directed to and will be fulfilled by the Lead Contact, Valerie M. Weaver. Modified plasmids and CAD drawings for 3D-printed parts can be made available upon request.

### Experimental Models and Subject Details

#### Cell Lines

Human embryonic stem cells (parental line H9, female) were obtained as a gift from the Laboratory of Susan Fisher at UCSF and maintained in a humidified incubator at 37°C with 5% CO_2_. T-mNeonGreen, H2B-mCherry, and shE-cadherin lines were all generated from H9s, as detailed in the subsequent sections of the Method Details.

T-mNeonGreen reporter cells were maintained on γ-irradiated primary mouse embryonic fibroblasts (PMEFs) in KSR media consisting of knockout-DMEM (Gibco) with 20% knockout serum replacement (Gibco), 2 mM L-glutamine (Gibco), 1 mM non-essential amino acids (Gibco), 1× antibiotic-antimycotic (Gibco), 100 μM β-mercaptoethanol, supplemented with 10 ng/ml bFGF (PeproTech), and passaged with collagenase type IV (Gibco) at 1 mg/ml in knockout-DMEM (Gibco). PMEFs, derived from CF-1 mice, were obtained as a gift from the Laboratory of Susan Fisher at UCSF and were cultured in a humidified hypoxic incubator at 37°C with 5% O_2_ and 5% CO_2_ on tissue culture plastic coated with 0.1% gelatin (Sigma) and in media consisting of DMEM, high glucose, with L-glutamine and sodium pyruvate (GenClone) supplemented with 10% fetal bovine serum (HyClone), 4 mM L-glutamine (Gibco), 1× antibiotic-antimycotic (Gibco), 10 ng/ml insulin (Roche), 20 ng/ml transferrin (Sigma), and 30 nM sodium selenite (VWR). PMEFs were γ-irradiated with a total dose of 40 Gy to induce mitotic arrest and frozen aliquots were stored in liquid nitrogen prior to being thawed and plated onto tissue culture plastic coated with 0.1% gelatin (Sigma) for passaging of T-mNeonGreen reporter cells.

H9, H2B-mCherry, and shE-cadherin lines were maintained in feeder-free conditions on tissue culture plastic coated with Matrigel (R&D Systems) in media consisting of 50% PMEF-conditioned KSR media and 50% complete Essential 8 media (E8; Gibco), supplemented with 10 ng/ml bFGF (PeproTech). PMEF-conditioned KSR media was generated by plating γ-irradiated PMEFs on tissue culture plastic coated with 0.1% gelatin (Sigma), feeding with KSR media supplemented with 4 ng/ml bFGF (PeproTech), and collecting and replacing the media every 24 hours for 10-14 days. Cells in feeder-free conditions were passaged with 0.5 mM EDTA (Fisher) in PBS and plated into media supplemented with 10 μM Y-27632 (ROCK inhibitor; Tocris) to promote survival. After 24 hours media was replaced with media lacking Y-27632.

The human embryonic kidney (HEK) 293T cell line, obtained as a gift from the Laboratory of Warren Pear at UPenn, was used for transfection and production of lentivirus, and was maintained in a humidified incubator at 37°C with 5% CO_2_. HEK 293T cells were cultured in DMEM, high glucose, with L-glutamine and sodium pyruvate (GenClone) supplemented with 10% fetal bovine serum (HyClone), 4 mM L-glutamine (Gibco), and 1× antibiotic-antimycotic (Gibco), and upon reaching 75% confluency were passaged with 0.05% trypsin-EDTA (Gibco).

All cell lines were routinely tested and confirmed to be negative for mycoplasma contamination. All experiments involving hESCs were approved by the University of California San Francisco Human Gamete, Embryo and Stem Cell Research Committee (UCSF GESCR) study number 11-05439.

### Method Details

#### Generation of T-mNeonGreen reporter

H9 hESCs expressing a C-terminal fusion of the T gene with the mNeonGreen fluorophore were prepared by CRISPR-Cas9 facilitated homology-directed repair (HDR). Guide RNAs (gRNAs) directing CRISPR-Cas9 double stranded cleavage near the end of the coding sequence of the human T gene were cloned into a modified derivative of pX330-U6-Chimeric_BB-CBh-hSpCas9 (Addgene; modification: Lakins et al., in preparation) and screened for activity using the Surveyor Assay (Transgenomics) following transient transfection in HEK 293T cells. The gRNA with target sequence 5’ - GCCTTGCTGCTTCACATGGA - 3’ demonstrated the best activity and was cloned into a second modified version of pX330 for use in H9 cells, which we call SpCas9 U6 gRNA (Lakins et al., in preparation). Briefly, this derivative features gRNA under control of the U6 promoter as in pX330, a T2A polyprotein of SpCas9 and the human RAD51 gene under transcriptional control of the tetracycline-regulated TetO heptamerized minimal CMV promoter, an expression cassette for the advanced reverse tetracycline transcriptional transactivator rtTA^s^-M22 (Urlinger et al., 2000), and the origin of replication from Epstein-Barr virus (OriP). For the targeting construct, approximately 1,000 base pairs upstream and downstream of the CRISPR-Cas9 cleavage site/T gene stop codon was prepared by Long Range PCR using a Vent-Tth Polymerase mix (Sigma; New England BioLabs; Cheng et al., 1994) and cloned into pBluescript II KS+ (Stratagene) modified by the addition of OriP. This targeting construct was further modified by silent mutation of the gRNA targeting sequence, removal of the natural T gene stop codon, and insertion of a 22 amino acid glycine/serine/alanine rich flexible linker N-terminal to mNeonGreen in frame with the T gene coding sequence. Following mNeonGreen, the construct included a Floxed expression cassette containing an mCherry fluorophore for assessing transfection efficiency and the puromycin resistance gene for selection of stably transfected cells.

H9 cells were co-transfected via electroporation with the described targeting construct, the SpCas9 U6 gRNA, and an *in vitro* transcribed capped, polyadenylated mRNA for the Epstein-Barr virus nuclear antigen 1 (EBNA-1) lacking the GA rich domain (Howden et al., 2006; targeting construct and SpCas9 U6 gRNA: Lakins et al., in preparation). Transfected H9s were re-plated on Matrigel-coated dishes in the presence of 10 µM Y-27632 (Tocris) to promote survival and 1 μg/ml of Doxycycline to induce the targeted CRISPR-Cas9 double-stranded break and subsequent homology-directed repair. Doxycycline was removed after 24 hours and cells were allowed to recover an additional 24 hours before selection with 0.25 μg/ml of puromycin. Following selection, individual surviving colonies were mechanically passaged into separate wells, expanded, and then screened for gene targeting via Long Range PCR of genomic DNA using one primer anchored in mNeonGreen and a second in the T gene upstream of the end of the 5’ homology arm. Positive clones were expanded and subsequently transiently transfected with a plasmid expressing an EGFP-Cre fusion to remove the puromycin selection cassette. Transfected cells were FACS-sorted for EGFP expression 24 hours later, re-plated in Matrigel-coated dishes in the presence of 10 µM Y-27632, and following outgrowth of individual colonies, mechanically passaged into separate wells and screened by PCR for loss of the expression cassette. These cells were then characterized for expression of nuclear mNeonGreen expression following BMP4 differentiation, and verified by the concordance of mNeonGreen expression and detection of the endogenous T gene by immunostaining.

#### Generation of H2B-mCherry

To generate the hESC line with fluorescent H2B-mCherry-labelled nuclei, an N-terminal fusion of human histone H2B to mCherry, under control of the human phosphoglycerate kinase promoter, was cloned into a transfer vector for transfection into HEK 293T cells, along with packaging and envelope vectors for self-inactivating transgenic lentivirus production, and subsequent transduction into H9s. Twenty-four hours prior to transfection, HEK 293T’s were plated at a density of 0.8 x 10^6^ cells into a 35 mm tissue culture plastic dish. The next day, HEK 293T’s were washed once with PBS very gently to remove serum, and media was replaced with Opti-MEM (Gibco). To begin transfection, 1 µg of total DNA consisting of 0.33 µg psPAX2 packaging vector (Addgene), 0.16 µg pMD2.G envelope vector (Addgene), and 0.55 µg H2B-mCherry transfer vector was mixed with Opti-MEM media to a total volume of 100 µL and incubated for 5 minutes at room temperature. Simultaneously, 3 µg of polyethylenimine (PEI; Sigma) was mixed with 100 µL Opti-MEM media and incubated for 5 minutes at room temperature. Following incubation, these two solutions were combined and incubated an additional 15 minutes at room temperature before being added to HEK 293T’s. Six hours following transfection, media was replaced for normal HEK 293T growth media. Forty-eight hours post-transfection, viral supernatant was collected, cell debris was removed with 2x 5-minute centrifugation at 500 x g, carefully collecting supernatant and discarding pellets each time, and polybrene (Sigma) was added to the final supernatant at 4 µg/ml. H9s plated on Matrigel-coated tissue culture plastic were immediately transduced by mixing viral supernatant 1:4 with E8 media (Gibco) supplemented with 10 ng/ml bFGF (PeproTech) and culturing cells in this media for 24 hours, after which the media was replaced with fresh media and cultured for an additional 48 hours. A pure population of cells expressing H2B-mCherry-labelled nuclei was obtained in two steps: first by enrichment via several rounds of mechanically picking morphologically undifferentiated colonies containing mCherry-positive cells and passaging as colony fragments onto γ-irradiated PMEFs, and then subsequently by FACS to purify. Single cells isolated by FACS were plated onto Matrigel-coated tissue culture plastic in E8 media (Gibco) supplemented with 10 ng/ml bFGF (PeproTech) and 10 μM Y27632 (Tocris) to promote survival. Fresh media was added every 24 hours and Y27632 was reduced to 5 μM, 2.5 μM, and 0 μM on successive days, after which self-supporting undifferentiated colonies were obtained.

#### Generation of shE-cadherin

We utilized the same inducible shE-cadherin hESC line from our previous work (Laralynne Przybyla et al., 2016). To generate the hESC line with inducible short hairpin RNA (shRNA) knockdown of E-cadherin, candidate shRNA hairpins were cloned into a transfer vector for transfection into HEK 293T cells along with packaging and envelope vectors for transgenic lentivirus production and subsequent transduction into H9s. The transfer vector consisted of a modified pLKO.1 neo plasmid (Addgene) with expression of the shRNA sequences under control of 3x copies of the lac operator, and contained a copy of the mNeonGreen fluorophore to assess transfection efficiency. The successful E-cadherin shRNA had the following sequence: 5’ - GAACGAGGCTAACGTCGTAAT - 3’. Transgenic lentivirus was produced in HEK 293T’s as described in the preceding section for generation of the H2B-mCherry line. H9s plated on Matrigel-coated tissue culture plastic were immediately transduced by mixing viral supernatant 1:4 with E8 media (Gibco) supplemented with 10 ng/ml bFGF (PeproTech) and culturing cells in this media for 24 hours, after which the media was replaced with fresh media and cultured for an additional 24 hours before selection with 200 µg/ml G-418 (Sigma). shE-cadherin cells were maintained with 200 µg/ml G-418 prior to seeding for experiments to prevent loss of inducible E-cadherin knockdown. E-cadherin knockdown was induced with 200 µM isopropyl-β-D-thiogalactoside (IPTG; Sigma).

#### Atomic Force Microscopy

Embryos were dissected from fertilized chicken eggs (Petaluma Farms) and cultured on top of filter paper to maintain tension across the blastoderm and vitelline membranes (Chapman et al., 2001) until Hamburger and Hamilton (HH) stage 3+ to 4, when the primitive streak is fully formed and mesodermal cells are actively ingressing (Hamburger and Hamilton, 1992; Voiculescu et al., 2014). Gastrulation-stage embryos were then mounted onto glass slides and placed on the stage of an MFP3D-BIO inverted optical AFM (Asylum Research) on a Nikon TE2000-U inverted microscope. Indentations were made using silicon nitride cantilevers with spring constants ranging from 0.05 to 0.07 N/m and borosilicate glass spherical tips 5 mm in diameter (Novascan Tech). The cantilevers were calibrated using the thermal oscillation method prior to each experiment. Embryos were indented at rates ranging from 0.75 to 1.25 mm/s with a maximum force of 4.5 nN. The Hertz model was applied to the force curves obtained from each indentation to calculate the elastic modulus (Young’s modulus, stiffness). Embryos were assumed to be incompressible; therefore, a Poisson’s ratio of 0.5 was used in the calculation of the elastic modulus.

#### Fabrication of Non-patterned Hydrogels

Polyacrylamide hydrogels were fabricated as described previously (Lakins et al., 2012; L. Przybyla et al., 2016). First #1 18 mm glass coverslips (Electron Microscopy Sciences) were modified with glutaraldehyde to promote attachment of polyacrylamide during polymerization. Coverslips were submerged in 0.2 M HCl (Fisher) overnight with gentle shaking, washed with ultrapure water, submerged in 0.1 M NaOH (Fisher) for 1 hour with gentle shaking, washed with ultrapure water, submerged in a 1:200 dilution of 3-aminopropyltrimethoxysilane (Acros Organics) in ultrapure water for 1 hour with gentle shaking, washed with ultrapure water, and finally submerged in a 1:140 dilution of 70% glutaraldehyde (Electron Microscopy Sciences) in PBS for 1 hour with gentle shaking, washed with ultrapure water and dried. Polyacrylamide solutions yielding the desired elastic moduli (E) were prepared and pipetted onto glutaraldehyde-modified coverslips with custom-made plastic spacers of approximately 100-200 µm thickness. For hydrogels of E = 2700 Pa, the polyacrylamide solution consisted of 7.5% acrylamide (Bio-Rad), 0.035% Bis-acrylamide (Bio-Rad), 1x PBS, 1% TEMED (Bio-Rad) and 1% potassium persulfate (Sigma). A Rain-X-coated (Rain-X) coverslip was placed atop each glutaraldehyde-modified coverslip with polyacrylamide solution to form a polyacrylamide “sandwich”, and these were incubated for 1 hour at room temperature to allow polymerization. The Rain-X-coated coverslips and plastic spacers were then removed and the glutaraldehyde-modified coverslips with attached polyacrylamide gels were placed into custom 3D-printed holders (CAD drawings available upon request) with rubber gaskets to form sealed wells with the polyacrylamide hydrogels at the bottom of each well.

Next, the surfaces of the polyacrylamide gels were functionalized with N-succinimidyl acrylamidohexanoic acid (N6), which was synthesized in the lab as previously described (Lakins et al., 2012), to enable ECM ligands to bind to the surface and promote cell attachment. A solution consisting of 50 mM HEPES (Fisher), pH 6.0, 0.01% Bis-acrylamide (Bio-Rad), 25% ethanol, 0.01% N6 (custom-synthesized), 0.002% Di(trimethylolpropane) tetraacrylate (Sigma), and 0.025% Irgacure D-2959 (Sigma) was prepared, pipetted into each well containing a polyacrylamide gel, and exposed for 5–10 minutes with a medium wavelength UV source (Spectroline EN-180, 306 nm peak). The gels were then washed 2x 10 minutes with ice-cold 25 mM HEPES (Fisher), pH 6.0 and 2x 10 minutes with ice-cold 0.9% NaCl (Fisher). Matrigel (R&D Systems) was diluted to a concentration of 250 μg/ml in ice-cold 100 mM HEPES (Fisher), 100 mM NaCl (Fisher), pH 8.0, added to each well containing a gel, and incubated at 4° C overnight with gentle rocking. The gels were then washed 5x 10 minutes with PBS and stored at 4° C in sterile conditions with PBS prior to beginning experiments.

#### Fabrication of Patterned Hydrogels

Patterned polyacrylamide hydrogels were fabricated as described previously (Muncie et al., 2019). Glutaraldehyde-modified coverslips were generated as described in the previous section for non-patterned hydrogels. Additional #1 18 mm coverslips (Electron Microscopy Services) were cleaned by overnight submersion in 1 M HCl (Fisher) with gentle rocking. Custom silicon wafers containing desired geometric patterns were generated using negative photoresist according to manufacturer’s instructions (SU-8; Kayaku Advanced Materials, Inc.). Polydimethylsiloxane (PDMS; Dow) stamps with desired patterns were then fabricated on these custom silicon wafers. Stencils were generated by placing these PDMS stamps onto flat slabs of PDMS, wicking a UV-curable adhesive NOA-74 (Norland Products, Inc.) between the stamps and the flat slabs, and curing the adhesive via 10-minute exposure with a medium wavelength UV source (Spectroline EN-180, 306 nm peak). Next, stencils were firmly pressed onto the acid-washed coverslips and incubated overnight at 4° C with 250 μg/ml Matrigel (R&D Systems) in ice-cold 100 mM HEPES (Fisher), 100 mM NaCl (Fisher), pH 8.0. Stencils were then removed from the patterned coverslips and the coverslips were washed by two sequential submersions in PBS and a single submersion into ultrapure water before being dried under nitrogen gas (Airgas).

Polyacrylamide gels were then fabricated as described in the preceding section, with polyacrylamide solution being pipetted onto a glutaraldehyde-modified coverslip with plastic spacer and a patterned coverslip being placed on top, with the Matrigel-patterned side in contact with the polyacrylamide. These polyacrylamide “sandwiches” were incubated for 1 hour at room temperature to allow polymerization, then submerged in PBS and incubated at room temperature with gentle rocking for an additional 2 hours. The patterned top coverslips were removed using a scalpel while the “sandwich” remained submerged in PBS, and then the glutaraldehyde-modified coverslips with attached polyacrylamide gels were assembled into custom 3D-printed holders (CAD drawings available upon request) with rubber gaskets to form sealed wells with the polyacrylamide hydrogels at the bottom of each well. The gels were then washed 5x 10 minutes with PBS and stored at 4° C in sterile conditions with PBS prior to beginning experiments.

#### Traction Force Microscopy

Polyacrylamide hydrogels were fabricated as described in the preceding sections and in previous work (Lakins et al., 2012; L. Przybyla et al., 2016), with 1 µm diameter fluorescent microspheres (Invitrogen) added to the polyacrylamide solution at a final concentration of 0.03% solids. Upon adding the polyacrylamide solution between two coverslips, the coverslip “sandwiches” were centrifuged in swing-buckets at 200 x *g* for 10 minutes with the Rain-X-coated coverslip at the bottom, and incubated in this configuration at room temperature for an additional 1 hour to allow for polymerization. The centrifugation step forced all the microspheres into a single XY-plane at what ultimately became the surface of the hydrogel upon removal of the Rain-X-coated coverslip.

hESC colonies were plated onto hydrogels as described in the following section. At timepoints for which traction force measurements were desired, images of the fluorescent microspheres (“stressed images”) were captured using a Nikon Eclipse TE200 U (Nikon) inverted microscope with a 10x objective, equipped with a motorized positioning stage (Prior Scientific HLD 117) and an ORCA Flash 4.0LT CMOS camera (Hamamatsu). hESCs were then lysed using 2% sodium dodecyl sulfate (Sigma) and images were captured of the fluorescent microspheres from the same regions of interest (“unstressed images”). For each region of interest, “stressed” and “unstressed” images were aligned using Fiji software plugin Linear Alignment with SIFT (Schindelin et al., 2012). Bead movements resulting from cell traction forces were determined using a Fiji software plugin for computing particle image velocimetry (PIV) measurements (Tseng et al., 2012), and these PIV measurements were converted to traction stresses using the Fiji software plugin FTTC (Tseng et al., 2012). MATLAB (MathWorks) was used to visualize the plots of traction stress magnitudes. The code generated to analyze and display traction force data is publicly available on GitHub (https://github.com/jmmuncie/TF_hESC).

Plots of average traction stress magnitudes for patterned colonies were generated by using brightfield images of the colonies to align individual traction stress magnitude maps. The traction stress maps were then cropped to a uniform size and magnitude values at each voxel were averaged across all the maps. The number of colonies used to generate average traction stress maps can be found in the figure captions. The normalized frequency distribution plot of average traction stresses for shE-cadherin control versus knockdown colonies was generated using the Kernel Density Estimation add-in for Excel (Microsoft) with an *h* value of 1.5.

#### Plating hESCs onto Hydrogels

Twenty-four hours prior to plating hESCs, knockout-DMEM (Gibco) was added to the hydrogels and incubated at 37° C in a humidified cell culture incubator with 5% CO_2_. Prior to plating on hydrogels, T-mNeonGreen reporter cells cultured on γ-irradiated PMEFs were passaged into secondary cultures in feeder-free conditions on tissue culture plastic coated with Matrigel (R&D Systems) and fed with PMEF-conditioned KSR media supplemented with 10 ng/ml bFGF (PeproTech). hESC lines that were maintained in feeder-free conditions (H9, H2B-mCherry, and shE-cadherin) were passaged to hydrogels directly. hESCs were released from Matrigel-coated plastic via 10-minute incubation at 37° C with 0.05% trypsin-EDTA (Gibco) supplemented with 10 μM Y-27632 (Tocris), and counted using a hemocytometer. Unconfined colonies of hESCs were generated on non-patterned hydrogels by placing custom 3D-printing plating guides (3 mm diameter; CAD drawings available upon request) onto the surface of the hydrogels and pipetting 20,000 cells through each opening of the plating guides. Guides were removed after incubation for 1 hour to ensure cell attachment. Patterned hESC colonies were generated by seeding 200,000 cells onto each of the patterned gels and very gently replacing the media after 3 hours to remove non-attached cells. T-mNeonGreen reporter cells were plated in PMEF-conditioned KSR media, while the other cell lines were plated in 50% PMEF-conditioned KSR, 50% E8 media (Gibco). For shE-cadherin experiments, IPTG (200 µM; Sigma) was added to the media one passage (72 hours) prior to seeding on hydrogels to induce shE-cadherin knockdown. All cell lines were plated in media supplemented with 10 ng/ml bFGF (PeproTech) and 10 μM Y-27632 (Tocris). The Y-27632 was diluted out of the media over the course of 72 hours to prevent dissociation of hESCs from the hydrogels: 24 hours after plating, media was replaced with media containing 5 μM Y-27632; 48 hours after plating, media was replaced with media lacking Y-27632.

#### BMP4 Differentiation

Differentiation was induced by removing maintenance media from hESC colonies on hydrogels and replacing it with Stemline II Hematopoietic Stem Cell Expansion Medium (Sigma) supplemented with 10 ng/ml bFGF (PeproTech) and 50 ng/ml BMP4 (PeproTech). hESC colonies were differentiated for the amount of time described for each experiment. For shE-cadherin experiments, IPTG (200 µM; Sigma) was added to the differentiation media in the knockdown condition to maintain shE-cadherin knockdown. For Src inhibition experiments, the Src-family kinase inhibitor (PP1; 10 μM; Sigma) or equal volume of vehicle (DMSO) was added to the standard differentiation media.

#### Time-lapse Imaging

hESC colonies seeded on polyacrylamide gels in custom 3D printed holders were loaded into a custom-built stage mount as previously described (L. Przybyla et al., 2016; CAD drawings for gel holders and stage mount available upon request), and differentiation was induced as described in the preceding section. The stage mount was sealed, enabling a low-pressure flow of mixed gas containing 5% CO_2_, 95% air (Airgas) to maintain the pH of standard differentiation media, and the mount was placed onto a motorized positioning stage (Prior Scientific HLD 117) attached to a Nikon Eclipse TE200 U (Nikon) inverted epifluorescent microscope. The microscope stage, condenser, and objectives were encased in a Plexiglas box and a forced air temperature feedback control (In Vivo Scientific) was used to maintain the temperature of the entire setup at 37° C. Images were captured using a 10x objective at specified timepoints.

“Gastrulation-like” nodes were manually counted in colonies of H2B-mCherry cells by identifying regions of increased signal of mCherry, indicative of increased cell density and ingression. “Gastrulation-like” nodes were manually counted in colonies of T-mNeonGreen reporter cells by identifying regions with detectable nuclear mNeonGreen expression above background levels. The NIS-Elements (Nikon) software was used to measure the size of gastrulation nodes for both cell lines.

Plots of average T-mNeonGreen expression at 30 hours of BMP4 stimulation for triangle and Pac-Man patterned colonies were generated by using brightfield images to align the colonies from each replicate and crop the images to a uniform size. Background signal was removed by subtracting the image of mNeonGreen expression at 20 hours of BMP4 stimulation (prior to detectable T expression above background) from the image of mNeonGreen expression at 30 hours of BMP4 stimulation for each colony. The intensity values at each pixel in the resulting images were then divided by the number of replicates for each experiment, and summed across all the replicates to generate average expression plots, which were visualized with MATLAB (MathWorks; script available at https://github.com/jmmuncie/TF_hESC).

#### Immunofluorescence Staining and Imaging

hESC colonies on polyacrylamide gels were fixed slowly in cold conditions to prevent detachment of cells from the hydrogels. Prior to fixation, hESC colonies on gels were placed on ice and gently rocked for 10 minutes. Media was removed, ice-cold 4% paraformaldehyde (Sigma) was carefully added, and the samples were fixed at 4° C overnight. Prior to fixation, samples for β-catenin-Y654 staining were briefly washed with a hot “TNS” solution of 0.03% Triton-X 100 (Sigma), 0.4% NaCl (Fisher) at 90° C, and shook vigorously for 30 seconds to remove cytosolic β-catenin but leave junction β-catenin intact, as described previously (Röper et al., 2018). A larger volume of ice-cold TNS was then immediately added to rapidly cool each sample, after which all the TNS was removed and replaced with ice-cold 4% paraformaldehyde and samples were fixed at 4° C overnight.

After fixation, all samples were washed 3x 10 minutes with PBS and then simultaneously blocked and permeabilized with a 1-hour incubation at room temperature in “IF Buffer” containing 0.1% bovine serum albumin (Fisher), 0.2% Triton-X 100 (Sigma), 0.05% Tween-20 (Sigma), 130 mM NaCl (Fisher), 13 mM Na_2_HPO_4_ (Fisher), 3.5 mM NaH_2_PO_4_ (Fisher), and 0.05% sodium azide (Sigma), supplemented with 10% goat serum (Sigma). Samples were then incubated overnight with primary antibodies diluted in IF Buffer plus 10% goat serum at 4° C with gentle rocking. The next day, samples were washed 3x 10 minutes with IF Buffer at room temperature and then incubated for 2 hours with secondary antibodies diluted in IF Buffer plus 10% goat serum at room temperature with gentle shaking. Samples were then washed 3x 10 minutes with IF Buffer, 1x 5 minutes with 0.5 µg/ml DAPI (Invitrogen) in PBS, and 1x 10 minutes with PBS. Samples were then removed from the custom gel holders, inverted onto #1 22 x 55 mm glass coverslips (VWF) and imaged. Epifluorescent images were captured using a Nikon Eclipse TE200 U (Nikon) inverted microscope with a 10x, 20x, or 60x objective and an ORCA Flash 4.0LT CMOS camera (Hamamatsu). Confocal images were captured using a Nikon Eclipse Ti inverted microscope (Nikon) equipped with a 60x objective, a CSU-X1 spinning disk confocal scanner (Yokogawa), and a Zyla sCMOS camera (Andor).

Primary antibodies used were: anti-T(brachyury) (RRID: AB_2200235, R&D Systems, 1:40), anti-E-cadherin (RRID: AB_2291471, CST, 1:200), anti-Slug (RRID: AB_2239535, CST, 1:400), anti-β-catenin-Y654 (RRID: AB_10623284, Sigma, 1:50), anti-β-catenin (RRID: AB_11127855, CST, rabbit, 1:200), anti-β-catenin (ECM Biosciences, mouse, 1:250), anti-phospho-Src Family (RRID: AB_10013641, CST, 1:100), anti-Oct-3/4 (RRID: AB_2167703, Santa Cruz Biotechnology, 1:100). Secondary antibodies used were: Alexa Fluor 488 goat anti-mouse IgG (RRID: AB_2576208, Abcam, 1:1000), Alexa Fluor 568 goat anti-mouse IgG (Abcam, 1:1000), Alexa Fluor 488 goat anti-rabbit IgG (RRID: AB_2630356, Abcam, 1:1000), Alexa Fluor 568 goat anti-rabbit IgG (RRID: AB_2576207, Abcam, 1:1000), Alexa Fluor 488 donkey anti-goat IgG (RRID: AB_2687506, Abcam, 1:1000). Note: when anti-T(brachyury) and Alexa Fluor 488 donkey anti-goat IgG antibodies were used, donkey serum was used in place of goat serum during all blocking and staining steps described above.

Images were analyzed in Fiji (Schindelin et al., 2012). Plots of relative mean fluorescence intensity were generated by cropping all images to equivalent sizes and measuring the raw integrated density (RawIntDen, sum of pixel values). The RawIntDen values were normalized by dividing the RawIntDen value for each replicate within an experiment by the maximum RawIntDen value measured within that experiment. These normalized values were then plotted. Line graphs of fluorescence intensity values were generated by manually drawing lines in the indicated regions of interest, measuring the pixel intensity values along those lines, and plotting the resulting values.

#### Quantitative PCR (qPCR)

Total RNA was isolated from either full colonies of hESCs or FACS-sorted populations of T-mNeonGreen reporters as indicated in the main text and figure captions using TRIzol (Invitrogen) according to the manufacturer’s protocol. cDNA was synthesized from RNA using M-MLV Reverse Transcriptase (BioChain) and Random Hexamers (Applied Biosystems) as primers. qPCR was performed in triplicates from 10 ng of RNA per reaction using PerfeCTa SYBR Green FastMix (Quantabio) on a Mastercycler RealPlex^2^ detection system (Eppendorf). All reactions for qPCR were performed using the following conditions: 95° C for 30 seconds followed by 40 cycles of a three-step reaction of denaturation at 95° C for 10 seconds, annealing at 65° C for 10 seconds, and further annealing at 68° C for 20 seconds to reduce the likelihood of non-specific products, with reads taken at the end of each 68° C step. At the end of each reaction, melting curves were generated to validate the quality of amplified products using the following conditions: 95° C for 15 seconds, 60° C for 15 seconds, ramp to 95° C in 10 minutes. The mean Ct values from triplicates were used to calculate the ΔCt values relative to GAPDH expression. The means of the ΔCt values from independent experiments were used to calculate mean fold change of expression using the 2^-ΔΔCt^ method. For each gene evaluated, the SEM of the ΔCt values was calculated and used to generate positive and negative error values in the 2^-ΔΔCt^ fold change space. Plots of qPCR data display bars representing the mean fold change ±SEM and individual points representing the fold change value for each experimental replicate relative to the mean. All primers used in this study are listed in Table S1.

#### Eyebrow Knife Experiment

Eyebrow knives were fabricated by first heating 5.75” borosilicate glass Pasteur pipets (Fisher) over a Bunsen burner and pulling to generate a narrower tip. Paraffin wax (Sigma) was used to attach a plucked human eyebrow hair to the narrow tip of the pipet. (Note: Variations in the thickness and density of eyebrow hair will affect the ability to effectively manipulate hESC colonies, therefore, different eyebrow hairs from multiple individuals should be tested when adopting this method). “Stressed” microsphere images were captured for triangle patterned colonies of T-mNeonGreen reporter cells, as described in the previous section, prior to being cut with the eyebrow knife. Next, a Nikon SMZ800 (Nikon) dissecting microscope was used to make precise cuts with an eyebrow knife through the corners of triangle patterned colonies and additional “stressed” microsphere images were captured following the eyebrow knife cuts. The colonies were differentiated with BMP4 and time-lapse imaging was conducted to monitor T-mNeonGreen expression, as described in previous sections. After 48 hours of differentiation, cells were lysed with 2% SDS (Sigma) and “unstressed” microsphere images were captured. Finally, the microsphere images were used to generate plots of traction force magnitude for the colonies before and after cutting with the eyebrow knife and compared to the spatiotemporal expression of T-mNeonGreen captured during the time-lapse imaging.

The plot of time to T-mNeonGreen expression was generated by analyzing the time-lapse imaging data at each corner of the triangle patterned colonies successfully cut with the eyebrow knife. Each corner was categorized as “cut” or “intact” and the timepoint at which nuclear expression of T-mNeonGreen was detectable above background for cells within each corner was recorded and plotted.

#### In Situ Hybridization via Hybridization Chain Reaction (ISH-HCR)

Third-generation ISH-HCR was performed as described previously (Choi et al., 2018). Samples were fixed slowly in cold conditions to prevent detachment of hESCs from the hydrogels. Prior to fixation, hESC colonies on gels were placed on ice and gently rocked for 10 minutes. Media was removed, ice-cold 4% paraformaldehyde (Sigma) was carefully added, and the samples were fixed at 4° C overnight. The next day, samples were washed 3x 10 minutes with diethyl pyrocarbonate (DEPC; Sigma) treated PBS with 0.1% Tween-20 (Sigma) to permeabilize, and then washed 1x 5 minutes with 5x SSC buffer with 0.1% Tween-20. Samples were incubated for 1 hour at 37° C in a humidified chamber in hybridization buffer consisting of 30% de-ionized formamide (Sigma), 5x SSC, 9 mM citric acid, pH 6.0 (Sigma), 0.1% Tween-20, 50 µg/ml heparin (Sigma), 1x Denhardt’s solution (Sigma), 10% dextran sulfate, avg M_w_ >500,000 (Sigma), and DEPC-treated ultrapure water. Samples were then hybridized overnight via incubation at 37° C in a humidified chamber with hybridization buffer plus 20 nM split initiator hybridization probes designed to target Wnt3a (Table S2) and 10 mM ribonucleoside vanadyl complexes (Sigma). The next day, samples were washed 5x 10 minutes at 37° C with no agitation using a buffer consisting of 30% formamide, 5x SSC, and 9 M citric acid, pH 6.0. Samples were further washed 3x 10 minutes at room temperature with gentle shaking using a buffer of 5x SSC, 0.1% Tween-20, and 50 µg/ml heparin. Samples were incubated for 30 minutes at room temperature in amplification buffer consisting of 5x SSC, 0.1% Tween-20, and 10% dextran sulfate, avg M_w_ >500,000. Samples were then incubated overnight at room temperature in amplification buffer plus 60 nM HCR3 amplification probes conjugated with Alexa Fluor 647 (Choi et al., 2018). The next day, samples were washed 5x 10 minutes at room temperature with gentle shaking in 5x SSC with 0.1% Tween-20. The third wash also contained 0.5 µg/ml DAPI (Invitrogen). Finally, samples were inverted onto #1 22 x 55 mm glass coverslips (VWF) and imaged using a Nikon Eclipse Ti inverted microscope (Nikon) equipped with a 60x objective, a CSU-X1 spinning disk confocal scanner (Yokogawa), and a Zyla sCMOS camera (Andor). Split initiator hybridization probe sequences targeting Wnt3a mRNA transcripts are listed in Table S2.

### Quantification and Statistical Analysis

Tests of significance for comparisons between two experimental groups were performed using the two-tailed unpaired Student’s t-test in Prism 6 software (GraphPad), except for the data presented in Figure panels 6D and 7D, for which one-tailed unpaired Student’s t-tests were used because previous results suggested the expected direction of change in gene expression levels. Tests of significance for comparisons between more than two experimental groups were performed using a one-way Analysis of Variance (ANOVA) test in Prism 6 software (GraphPad). Differences were determined to be statistically significant at p < 0.05, and statistical significance was denoted by asterisks in the figure panels, with * = p < 0.05, ** = p < 0.01, and *** = p < 0.001. Differences that were determined to not be statistically significant were either denoted “n.s.” or the p-value was noted above the data. For immunofluorescent image data, the figure panels illustrate representative results from n = 3 independent experiments where a minimum of three replicate hESC colonies were imaged for each condition in each experiment. For qPCR data n = the number of independent experiments, with hESCs pooled from at least three different hydrogels for each condition in each experiment. All other definitions of n can be found in the figure captions. Statistical parameters plotted in each figure are also described in the figure captions.

### Data and Code Availability

The code generated during this study for analysis and visualization of traction force microscopy data is publicly available on GitHub: https://github.com/jmmuncie/TF_hESC.

### Key Resources Table

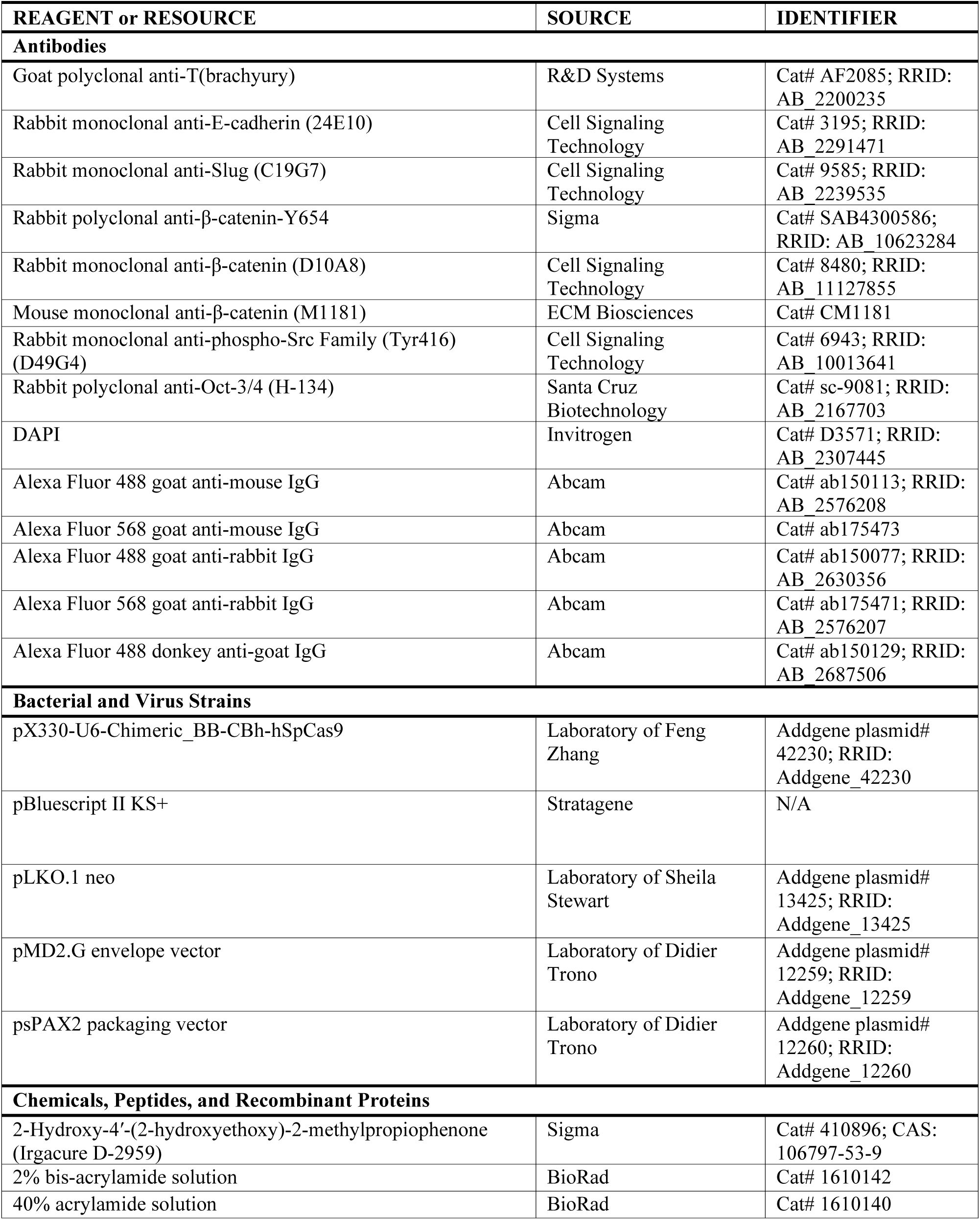
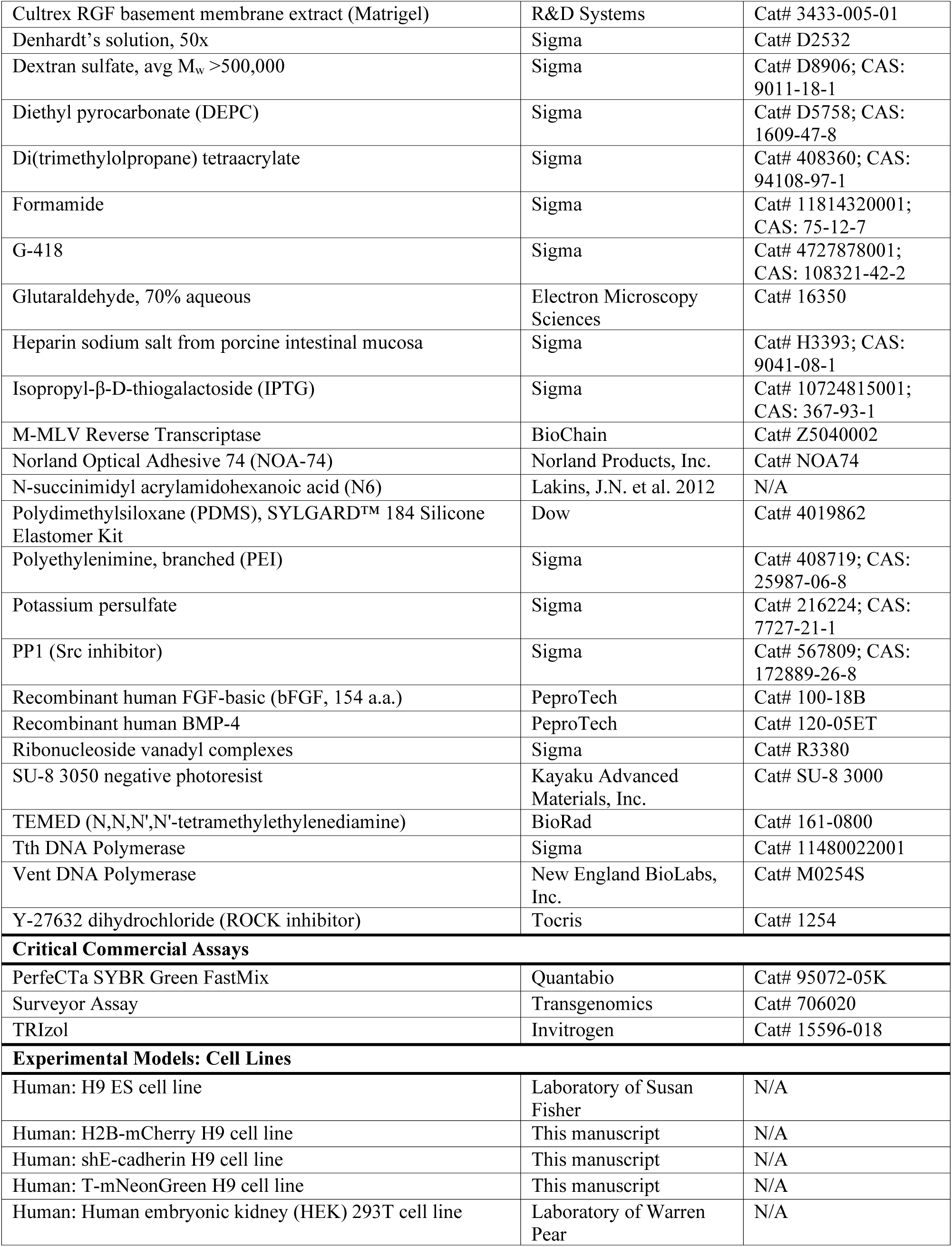
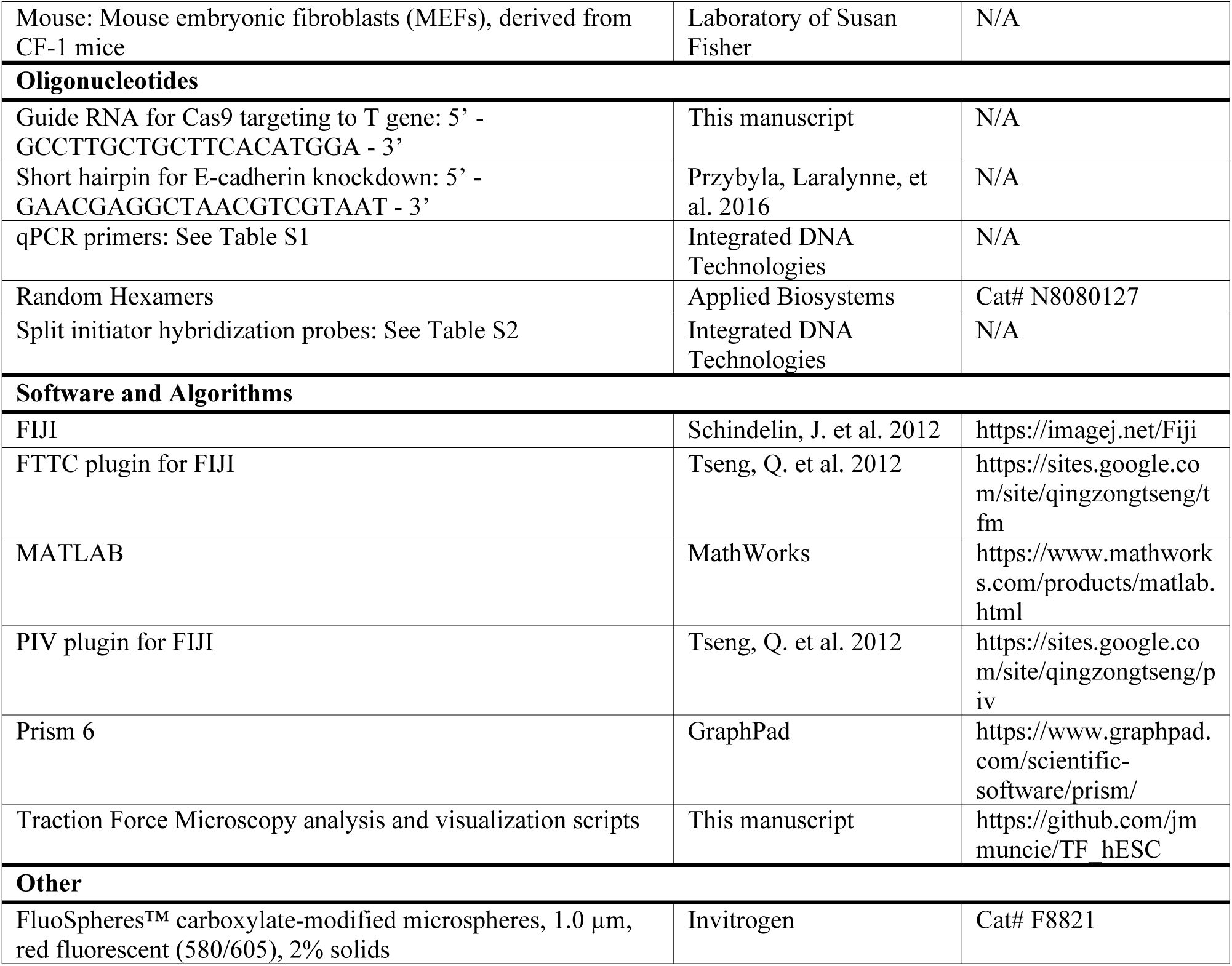

