## Supplemental Data for "Mechanics regulate human embryonic stem cell self-organization to specify mesoderm"

**Figure S1, related to Figure 1**

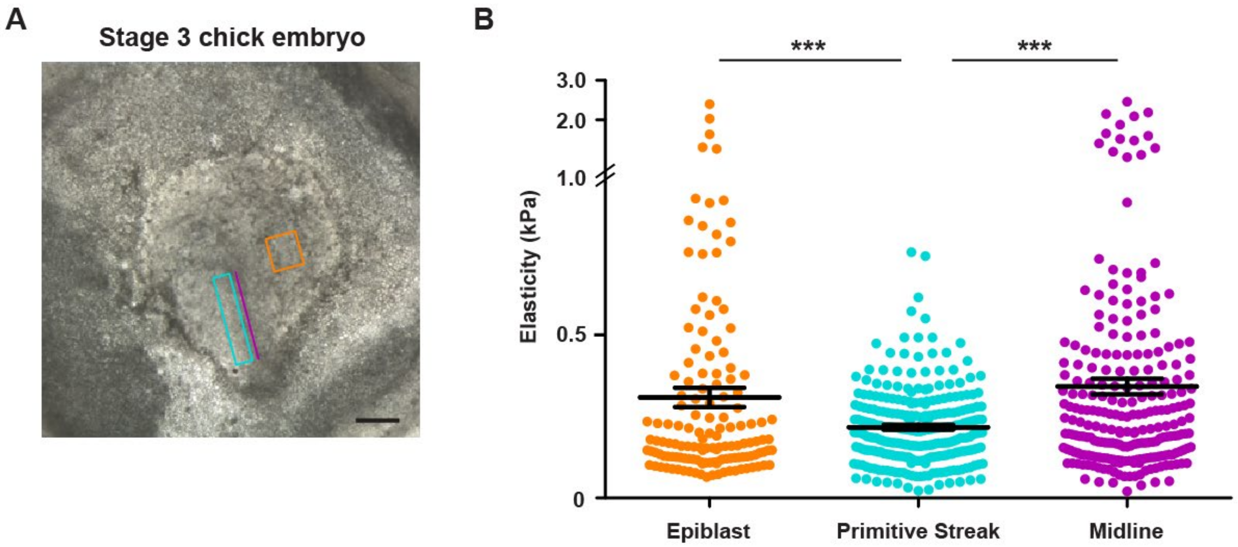


**Figure S1, related to Figure 1: Compliance of the extracellular microenvironment of gastrulation-stage chick embryos. A)** Representative brightfield image of an HH stage 3 chicken embryo extracted and prepared for atomic force microscopy measurements. Colored rectangles and line on the brightfield image indicate regions within the embryo where measurements were taken, corresponding to the data presented in panel B. Scale bar = 500 µm. **B)** Atomic force microscopy measurements of the elasticity of the extracellular microenvironment of HH stage 3 chicken embryos, grouped by regions of the embryo within which the measurements were taken. Each data point represents a single measurement of elasticity, with data collected from embryos prepared on four separate days, with at least three embryos measured each day. Epiblast n = 140, Primitive Streak n = 260, Midline n = 236. Bars represent mean ± SEM. ***$p < 0.001$. kPa = kilopascals.

**Figure S2, related to Figure 2**

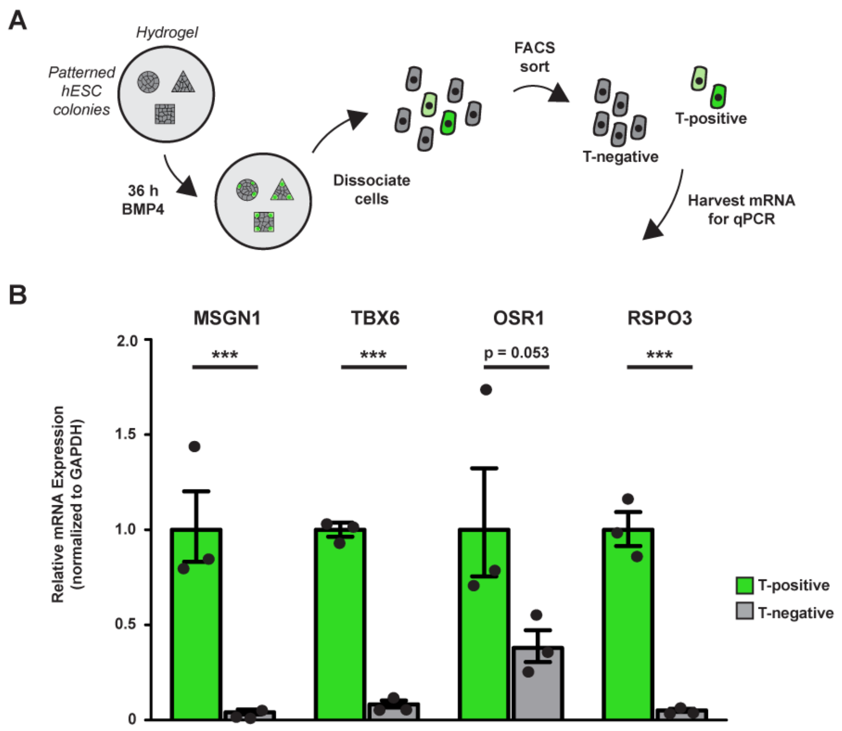


**Figure S2, related to Figure 2: T-reporter is capable of driving expression of direct transcriptional targets of T(brachyury). A)** Cartoon of protocol used to compare gene expression of direct T targets between T-mNeonGreen-positive and T-mNeonGreen-negative cells following 36 h of BMP4 stimulation. **B)** Bar graphs showing mRNA levels of direct transcriptional targets of T(brachyury) in T-positive and T-negative cells isolated from geometrically-confined hESC colonies on compliant hydrogels following 36 h of BMP4 stimulation. Bars represent mean fold change ± SEM from three independent experiments. ***$p < 0.001$. MSGN1 = Mesogenin 1, TBX6 = T-Box Transcription Factor 6, OSR1 = Odd-Skipped Related Transcription Factor 1, RSPO3 = R-Spondin 3.

**Figure S3, related to Figure 3**

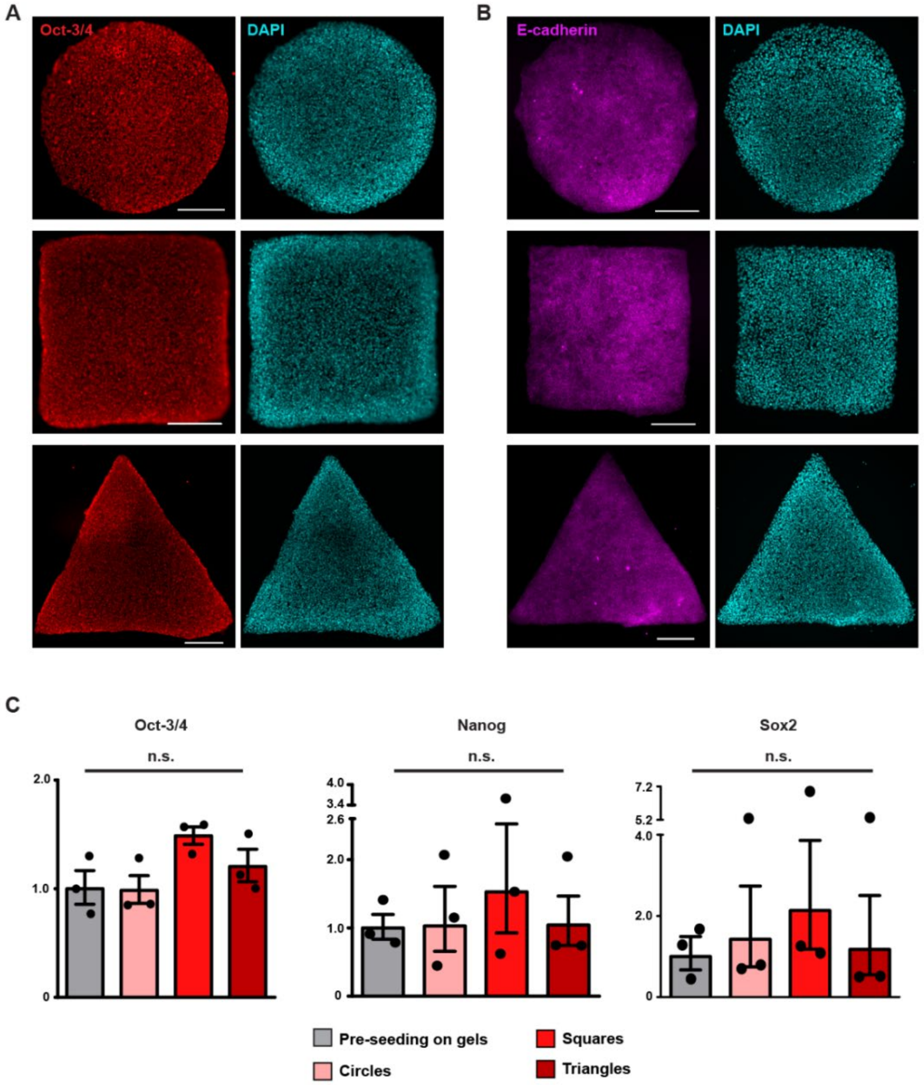


**Figure S3, related to Figure 3: H9s seeded on patterned gels remain pluripotent in maintenance conditions. A)** Representative immunofluorescent images of Oct-3/4 (Alexa568; red; left) and nuclei (DAPI; cyan; right) in geometrically-colonies of hESCs on compliant hydrogels in maintenance conditions. Scale bars = 250 µm. **B)** Representative immunofluorescent images of E-cadherin (Alexa568; magenta; left) and nuclei (DAPI; cyan; right) in geometrically-confined colonies of hESCs on compliant hydrogels in maintenance conditions. Scale bars = 250 µm. **C)** Bar graphs showing mRNA levels of pluripotency genes in geometrically-confined colonies of hESCs on compliant hydrogels in maintenance conditions, relative to levels prior to seeding on hydrogels. Bars represent mean fold change ± SEM from three independent experiments. n.s. = not significant.

**Figure S4, related to Figure 7**

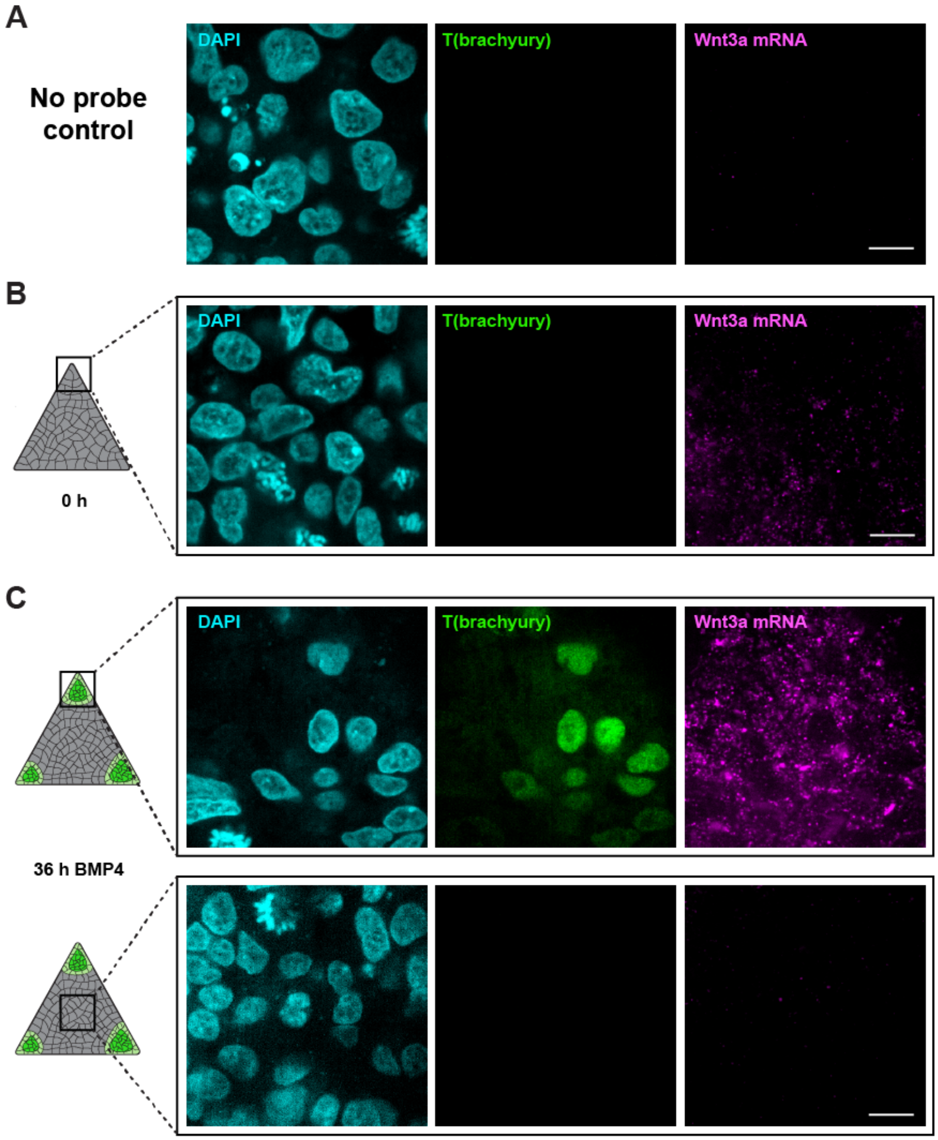


**Figure S4, related to Figure 7: Wnt3a in situ hybridization via hybridization chain reaction before and after BMP4 treatment. A)** Representative immunofluorescent images of nuclei (DAPI; cyan; left), immune-detected T(brachyury) protein (mNeonGreen; green; center) and *in-situ*-detected Wnt3a mRNA (Alexa647; magenta; right) in geometrically-confined triangle hESC colonies with no split initiator probes targeting Wnt3a prior to amplification with fluorescent probes. **B)** Same as in panel A, but including split initiator probes in geometrically-confined triangle hESC colonies prior to stimulation with BMP4. **C)** Same as in panel B, but following 36 h of BMP4 stimulation. The rectangles on the colony cartoons indicate the regions of the colonies where images were taken. All scale bars = 10 µm.

**Table S1, related to STAR Methods, Quantitative PCR (qPCR): Primers used for pPCR.**

| **Gene** | **Forward Primer Sequence, 5' to 3'** | **Reverse Primer Sequence, 5' to 3'** |
|---|---|---|
| GAPDH | CAGCCTCAAGATCATCAGCA | TGTGGTCATGAGTCCTTCCA |
| T(brachyury) | CAGCAAAGTCAAGCTCACCA | TGGACCCCCAACTCTCACTA |
| Goosecoid (GSC) | TCTCAACCAGCTGCACTGTC | CGTTCTCCGACTCCTCTGAT |
| Snai2 | TTGTGTTTGCAAGATCTGCGG | TGCAAATGCTCTGTTGCAGT |
| Sox2 | AGGATAAGTACACGCTGCCC | TAACTGTCCATGCGCTGGTT |
| E-cadherin | CGGCCTGAAGTGACTCGTA | GCCGCTTTCAGATTTTCATC |
| Wnt3a | GCCCCACTCGGATACTTCTT | GAGGAATACTGTGGCCCAAC |
| Wnt8a | TGTGATGGGTCAAACAATGG | TCCTTCCCCTTCTCCAAACT |
| Wnt4 | CCCTCATGAACCTCCACAAC | ACCTCACAGGAGCCTGACAC |
| Oct-3/4 | AGTGAGAGGCAACCTGGAGA | ACACTCGGACCACATCCTTC |
| Nanog | AGATGCCTCACACGGAGACT | AAGTGGGTTGTTTGCCTTTG |
| Msgn1 | TGTTGGACCCACCAGAACAC | TTGCAAAGGATGAGCCTCCC |
| Tbx6 | GAACCGGGAGCTATGGAAGG | AGAAACAAGTAGCGGGCCTC |
| Osr1 | TCCCTGGTTCCCTCATGTCA | CGGATCTTCTTGCGTTGCTG |
| Rspo3 | ACTTGCGACTGATTTCTTGGC | TCCTTGGCAGCCTTGACTAA |

**Table S2, related to STAR Methods, In Situ Hybridization via Hybridization Chain Reaction (ISH-HCR): Split initiator hybridization probe sequences used for Wnt3a ISH-HCR.**

| Probe | Sequence, 5' to 3' |
|---|---|
| Wnt3a_1a | AAAGTCTAATCCGTCCCT TT AGTAAGAAGTATCCGAGTGGGGCCA |
| Wnt3a_1b | CCCAGAGCCTGCTTCAGGCTGCAGA TT GCCTCTATATCTCCACTC |
| Wnt3a_2a | AAAGTCTAATCCGTCCCT TT CAGCGACCACCAGATCGGGTAGCTG |
| Wnt3a_2b | CAGGGAGGAATACTGTGGCCCAACA TT GCCTCTATATCTCCACTC |
| Wnt3a_3a | AAAGTCTAATCCGTCCCT TT TAGTTCCTGCAGAAGCGGAGCTGCT |
| Wnt3a_3b | TCGGCCACGCTGGGCATGATCTCCA TT GCCTCTATATCTCCACTC |
| Wnt3a_4a | AAAGTCTAATCCGTCCCT TT CCGAAGATGGCCAGGCTGTCGTGGA |
| Wnt3a_4b | TCCCTGGTAGCTTTGTCCAGCACGG TT GCCTCTATATCTCCACTC |
| Wnt3a_5a | AAAGTCTAATCCGTCCCT TT CTGAGGCAATGGCGTGGACAAAGGC |
| Wnt3a_5b | AGCGTGTCACTGCAAAGGCCACACC TT GCCTCTATATCTCCACTC |
| Wnt3a_6a | AAAGTCTAATCCGTCCCT TT ACTTCCAGCCCTTGCCTGGTGAGCC |
| Wnt3a_6b | ACTCGATGTCCTCGCTACAGCCACC TT GCCTCTATATCTCCACTC |
| Wnt3a_7a | AAAGTCTAATCCGTCCCT TT GGTTGCGACCACCAGCATGTCTTCA |
| Wnt3a_7b | AGGAAGTCACCGATGGCGCGGAAGT TT GCCTCTATATCTCCACTC |
| Wnt3a_8a | AAAGTCTAATCCGTCCCT TT ACCTTGAAGTAGGTGTAGCGCGGCC |
| Wnt3a_8b | TAGTAGACCAGGTCGCGCTCCGTGG TT GCCTCTATATCTCCACTC |
| Wnt3a_9a | AAAGTCTAATCCGTCCCT TT AGTGGAACACGCAGCGGCACTTCTC |
| Wnt3a_9b | ACTCCTGGCAGCTGACGTAGCAGCA TT GCCTCTATATCTCCACTC |
| Wnt3a_10a | AAAGTCTAATCCGTCCCT TT CAGGGAAAAGCCCACCCTCAGGCAG |
| Wnt3a_10b | CCGTTTAGGTGGGAGTCCTGCTCCA TT GCCTCTATATCTCCACTC |
| Wnt3a_11a | AAAGTCTAATCCGTCCCT TT AGCCCTGCCTTCAGGTAGGAGTTCT |
| Wnt3a_11b | AGAGAGGAGACACTAGCTCCAGGGA TT GCCTCTATATCTCCACTC |
| Wnt3a_12a | AAAGTCTAATCCGTCCCT TT TGGAGCTCCGCCTCATTCAGGAGCA |
| Wnt3a_12b | AAGCCAACGCAGAGCCCCTCCCCAT TT GCCTCTATATCTCCACTC |
| Wnt3a_13a | AAAGTCTAATCCGTCCCT TT GCCCAATCTGTAGCCCCGCCTCTGT |
| Wnt3a_13b | AGCCCTGTCCCACCCAAGAGAAGCC TT GCCTCTATATCTCCACTC |
| Wnt3a_14a | AAAGTCTAATCCGTCCCT TT ACCCAGAGCCACGCCCTTACTGGGA |
| Wnt3a_14b | GGTAGAAGCCTACCTAGTGCCCCGC TT GCCTCTATATCTCCACTC |
| Wnt3a_15a | AAAGTCTAATCCGTCCCT TT ATAAAACCCCACTCCTAAGGAGGCG |
| Wnt3a_15b | CCCATCCAGGAAGAAGCCTCATCCA TT GCCTCTATATCTCCACTC |
| Wnt3a_16a | AAAGTCTAATCCGTCCCT TT AAGGAGCCTATGCAGGCCACGCCCA |
| Wnt3a_16b | TGGTCCCAGAGAAGCCCCACCCACA TT GCCTCTATATCTCCACTC |
| Wnt3a_17a | AAAGTCTAATCCGTCCCT TT GAAGAGTCCCACCCGCGGAGAGAAG |
| Wnt3a_17b | GCCTTAATCAGGAGGGCGGTTCCCA TT GCCTCTATATCTCCACTC |
| Wnt3a_18a | AAAGTCTAATCCGTCCCT TT TCTGGAGCCGGGATTCCTGCAGAAG |
| Wnt3a_18b | AGGTGGCTGGTGGGCTGAATTTCCT TT GCCTCTATATCTCCACTC |
| Wnt3a_19a | AAAGTCTAATCCGTCCCT TT ATGGAACCTTACAGGGGGTTGGGGA |
| Wnt3a_19b | AACCTTCCCAGCTCGACGCAGGGGT TT GCCTCTATATCTCCACTC |
| Wnt3a_20a | AAAGTCTAATCCGTCCCT TT TGGGTGGTCAAACCCCAGGCTGAGG |
| Wnt3a_20b | TTTCCCCAGGTAGGGCCCCTGGTCA TT GCCTCTATATCTCCACTC |
